# An efficient opal-suppressor tryptophanyl pair creates new routes for simultaneously incorporating up to three distinct noncanonical amino acids into proteins in mammalian cells

**DOI:** 10.1101/2022.08.02.502539

**Authors:** Arianna O. Osgood, Yunan Zheng, Soumya Jyoti Singha Roy, Conor Loynd, Delilah Jewel, Abhishek Chatterjee

## Abstract

The site-specific incorporation of multiple distinct noncanonical amino acids (ncAAs) into proteins in mammalian cells is an emergent technology with much potential. For each different ncAA to be incorporated, this technology requires a distinct orthogonal aminoacyl-tRNA synthetase (aaRS)/tRNA pair that recognizes a distinct nonsense codon. The aaRS/tRNA pairs currently available for ncAA mutagenesis in eukaryotes are all traditionally used to decode the TAG nonsense codon. Unfortunately, these pairs suppress the other two nonsense codons, TGA or TAA, at a significantly lower level, compromising the scope of multi-ncAA mutagenesis. Here we report that the bacteria-derived tryptophanyl (EcTrp) pair is an excellent TGA-suppressor in mammalian cells. Additionally, we show that this pair does not cross-react with any of the three previously established aaRS/tRNA pairs. Consequently, the TGA-suppressing EcTrp pair can be combined with TAG-suppressing pyrrolysyl (archaeal), tyrosyl (bacterial), or leucyl (bacterial) pairs to develop three new routes for dual-ncAA incorporation in mammalian cells. We show that all three platforms enable site-specific incorporation of two distinct ncAAs into proteins – including a full-length humanized antibody – with excellent fidelity and good efficiency. Finally, we combined the EcTrp pair with the bacterial Tyr pair and the archaeal pyrrolysyl pair to site-specifically incorporate different combinations of three distinct ncAAs into a reporter protein in mammalian cells.

## Introduction

The ability to introduce multiple distinct non-natural functionalities at precise locations within a protein holds potential for numerous applications to probe and engineer protein structure and function.^1–3^ An attractive strategy to achieve this involves co-translational incorporation of noncanonical amino acids (ncAAs) into proteins through expanding the genetic code of the host cell.^4–9^ This method offers numerous advantages, including the flexibility of incorporating the desired functionality at virtually any site within the protein, the relatively small size of ncAAs compared to other genetically encoded tags, and the large available toolbox of genetically encoded ncAAs.

An ncAA of interest is incorporated into a protein in response to a nonsense or frameshift codon, using an engineered aminoacyl synthetase (aaRS)/tRNA pair that does not cross-react with its counterparts from the host cell.^4–6^ To simultaneously incorporate multiple different ncAAs into a protein, each ncAA must be assigned to a distinct aaRS/tRNA pair that decodes distinct nonsense/frameshift codons.^4,6,10^ These different aaRS/tRNA pairs should not only be orthogonal to their host counterparts, but also to each other (i.e., mutually orthogonal). In bacterial expression systems, multi-ncAA mutagenesis technology has advanced rapidly over the last decade;^11–20^ successful incorporation of up to four distinct ncAAs into one protein has recently been reported.^21^ In contrast, similar applications in mammalian cells have remained underdeveloped.

Several different aaRS/tRNA pairs have been developed for ncAA mutagenesis in mammalian cells, including tyrosyl (EcTyr)^22,23^ and leucyl (EcLeu)^24,25^ pairs derived from *E. coli*, and different pyrrolysyl (Pyl) pairs derived from archaea.^16^ The following combinations of these pairs were also found to be mutually orthogonal, enabling site-specific incorporation of up to two distinct ncAAs into proteins in mammalian cells: Pyl+EcTyr and Pyl+EcLeu,^10,25–27^ as well as combinations of mutually orthogonal Pyl pairs.^28–30^ Unfortunately, while these available pairs are good suppressors of the TAG nonsense codon, their efficiency of decoding the TGA and TAA nonsense codons are significantly worse. Development of new mutually orthogonal aaRS/tRNA pairs, capable of efficiently decoding nonsense codons other than TAG, would significantly expand the scope of multi-ncAA mutagenesis technology in mammalian cells. Here, we show that the recently developed *E. coli* derived tryptophanyl-tRNA synthetase (EcTrpRS)/tRNA pair^31,32^ is an excellent TGA suppressor in mammalian cells. We further demonstrate that the TGA-suppressing EcTrp pair is orthogonal to TAG-suppressing EcTyr, EcLeu, and Pyl pairs, and these combinations can indeed be used to incorporate two distinct ncAAs into proteins in mammalian cells, including two mutually compatible bioorthogonal handles for concurrent protein labeling at two different sites. Finally, we combined EcTrp(TGA), EcTyr(TAG), and Pyl(TAA) pairs to simultaneously incorporate different combinations of three distinct ncAAs, including three mutually compatible bioconjugation handles,^18^ into a protein in mammalian cells. Despite the reassignment of all three stop codons, we ensured homogeneity at the C-terminus using a self-cleaving protein tag.

## Results and Discussion

### EcTrpRS/tRNA^EcTrp^ is an active TGA suppressor in mammalian cells

Site-specific incorporation of two distinct ncAAs relies on the use of two unique ‘blank’ codons. All aaRS/tRNA pairs previously developed for ncAA incorporation in mammalian cells were developed as TAG suppressors, and these are significantly more efficient at suppressing TAG than TGA or TAA.^10^ We recently developed the EcTrpRS/tRNA pair for ncAA incorporation in both eukaryotes and an altered translational machinery *E. coli* strain (ATMW), which exhibited efficient TGA suppression in ATMW *E. coli*.^31,32^ We wondered if its TGA suppression activity would also translate to mammalian cells. We compared the TAG and TGA suppression efficiency of four aaRS/tRNA pairs (EcTrp, EcTyr, EcLeu, and Pyl) by co-transfecting three separate plasmids containing the aaRS, its cognate TAG or TGA suppressing tRNA, and an EGFP reporter harboring either a TAG or TGA stop codon (Figure S1). The relative expression levels of the EGFP reporter were used to quantify the suppression efficiency. For this experiment, we used the following synthetases and ncAAs: an *M. barkeri* PylRS with N^ε^-boc-lysine (Bock, **1**; Fig. 1),^10^ a polyspecific EcTyrRS with O-methyltyrosine (OMeY, **2**; Figure 1),^10^ a polyspecific EcLeuRS with Cy5Az (**5**; Figure 1),^25^ and a polyspecific EcTrpRS with 5-hydroxytryptophan (5HTP, **4**; Figure 1).^31^ As reported before, Pyl, EcTyr, and EcLeu pairs suppressed TAG significantly more efficiently relative to TGA. In contrast, the EcTrp pair exhibited significantly higher TGA suppression efficiency relative to the other pairs; in fact, its TGA suppression efficiency was somewhat higher than TAG (Figure S1).

**Figure 1.**
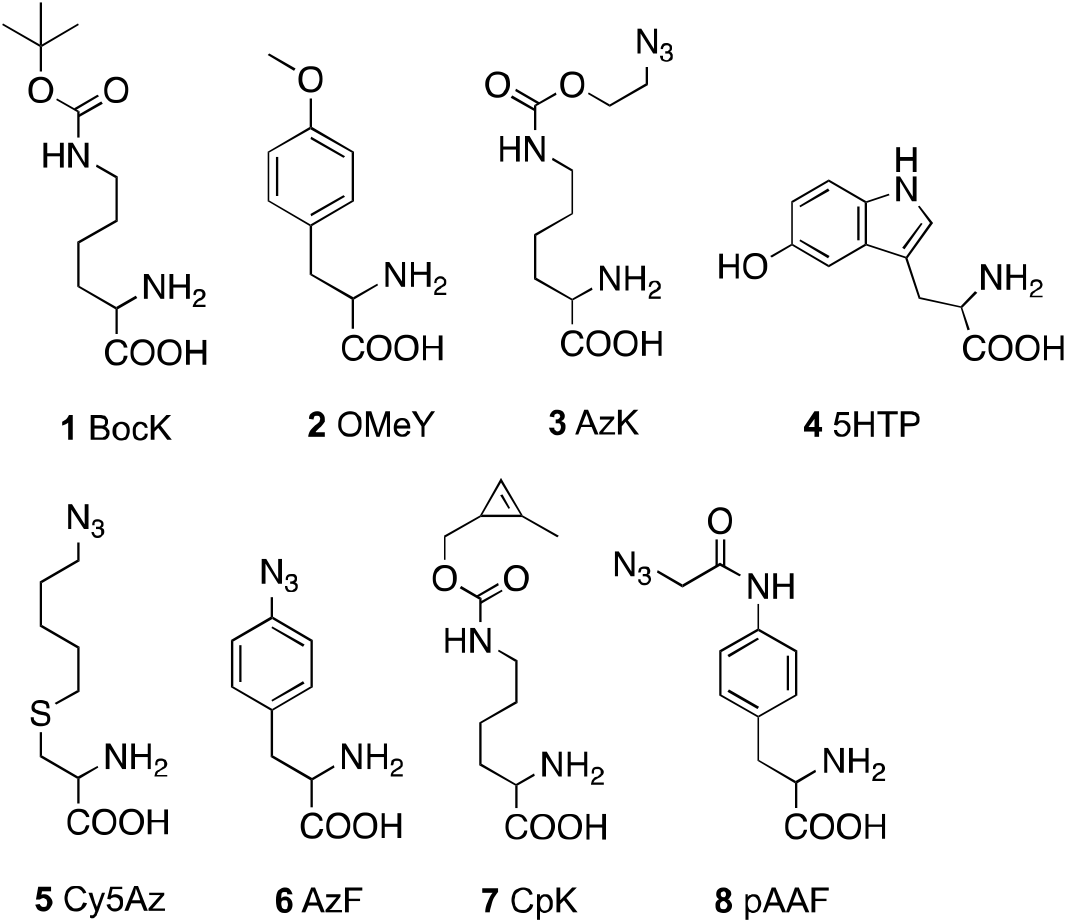
Structures of ncAAs used in this work.

### The EcTrpRS/tRNA^EcTrp^_UCA_ pair is orthogonal to three other mammalian suppressor pairs

High TGA suppression efficiency exhibited by the EcTrp pair makes it a good candidate for developing new dual-ncAA mutagenesis platforms in mammalian cells, provided it can be concurrently used with another TAG-suppressing pair without cross-reaction. In previous work, we have systematically evaluated potential cross-reactivity between the Pyl, EcLeu, and EcTyr pairs in mammalian cells, and showed that the Pyl is orthogonal to both EcLeu and EcTyr, but significant cross-reactivity exists between EcTyrRS and the tRNA_CUA_^EcLeu^.^10^ To explore if these three pairs can be combined with the opal suppressing EcTrp pair, we used the aforementioned three-plasmid transfection assay to probe potential cross-reactivities. Each aaRS was co-transfected with the four different TAG or TGA suppressing tRNAs along with an EGFP reporter harboring the appropriate nonsense codon (Figure 2). We observed robust reporter expression only when each aaRS was combined with its cognate suppressor tRNA, either for TAG (Figure 2B) or TGA (Figure 2C). The only exception was caused by the previously reported crossreactivity between EcTyrRS and tRNA^EcLeu^. These observations show that the TGA-suppressing EcTrp pair does not cross-react with the other three aaRS/tRNA pairs, and it should be possible to combine them to develop multiple new platforms for dual-ncAA incorporation in mammalian cells.

**Figure 2.**
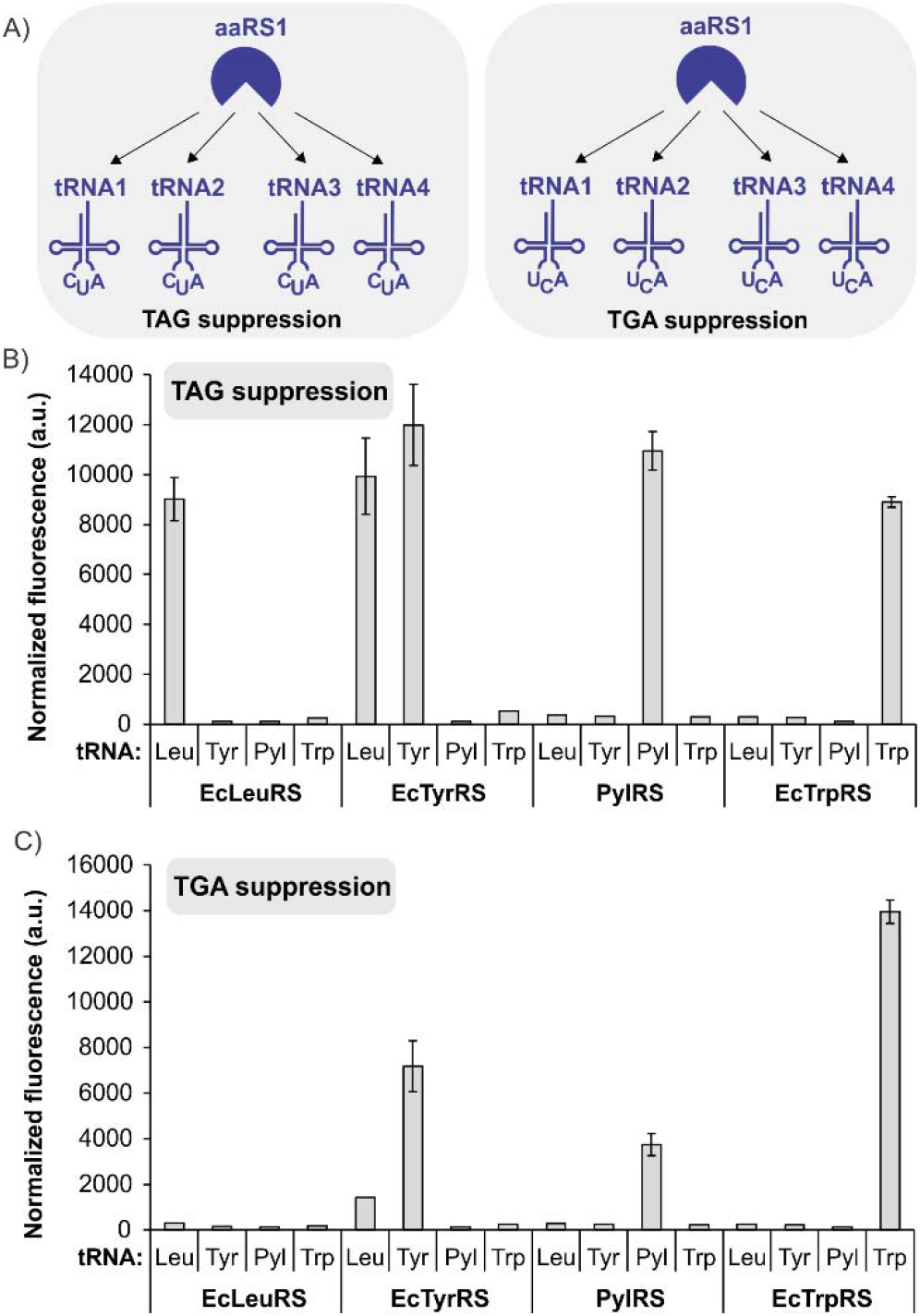
EcTrpRS/tRNA is orthogonal to the other aaRS/tRNA pairs in mammalian cells. A) The scheme of the experiment. Each aaRS was separately co-transfected into HEK293T cells with four different suppressor tRNAs (amber or opal suppressor), along with the appropriate EGFP-reporter and an appropriate ncAA. TAG and TGA suppression data are shown in panel B and C, respectively. Each aaRS only facilitates reporter expression in the presence of its cognate tRNA, except for the known cross-reaction between EcTyrRS and tRNA^EcLeu^. The EcTrp pair shows high opal suppression efficiency, at par with the amber suppression efficiency of other pairs.

### Three new dual suppression platforms using EcTrpRS/tRNA_UCA_^EcTrp^

To explore new dual suppression systems leveraging the opal suppressing EcTrp pair, we next focused on developing suitable plasmid systems based on previous successful designs.^10,25^ Expression vectors for each system included an expression cassette of the appropriate aaRS and multiple copies of the corresponding suppressor tRNA. For each dual suppression combination, one of the expression vectors also encoded an EGFP harboring a TAG and a TGA codon at permissive sites. Plasmids containing EcTyr_TAG_+EcTrp_TGA_, EcLeu_TAG_+EcTrp_TGA_, or Pyl_TAG_+EcTrp_TGA_ were co-transfected into HEK293T cells and reporter expression was evaluated by measuring fluorescence. Gratifyingly, all three dual suppression systems tested showed successful reporter expression in the presence of two appropriate ncAA substrates (Figure 3). Although reporter expression significantly diminished in the absence of either of the two ncAA substrates, noticeable expression was observed when 5HTP alone was omitted. This is likely due to the ability of the engineered EcTrpRS mutant to charge tryptophan at low levels in the absence of its preferred substrate 5HTP.^18,31,32^ However, we have previously shown that 5HTP is exclusively incorporated, when it is supplemented in the medium.^18,31,32^ Each of the three engineered synthetases used here are highly polyspecific (the ability to recognize multiple structurally similar ncAAs). Taking advantage of their polyspecificity, we were further able to show dual suppression using different ncAA combinations: OMeY+5HTP and AzF+5HTP for EcTyr_TAG_+EcTrp_TGA_ (Figure 3B); Cy5Az+5HTP for EcLeu_TAG_+EcTrp_TGA_ (Figure 3C); BocK+5HTP and AzK+5HTP for Pyl_TAG_+EcTrp_TGA_ (Figure 3D). Successful incorporation of the ncAAs was further confirmed by isolating mutant proteins from each dual suppression system using the C-terminal polyhistidine tag followed by SDS-PAGE (Figure S2) and whole protein ESI-MS analysis (Figure 3E).

**Figure 3.**
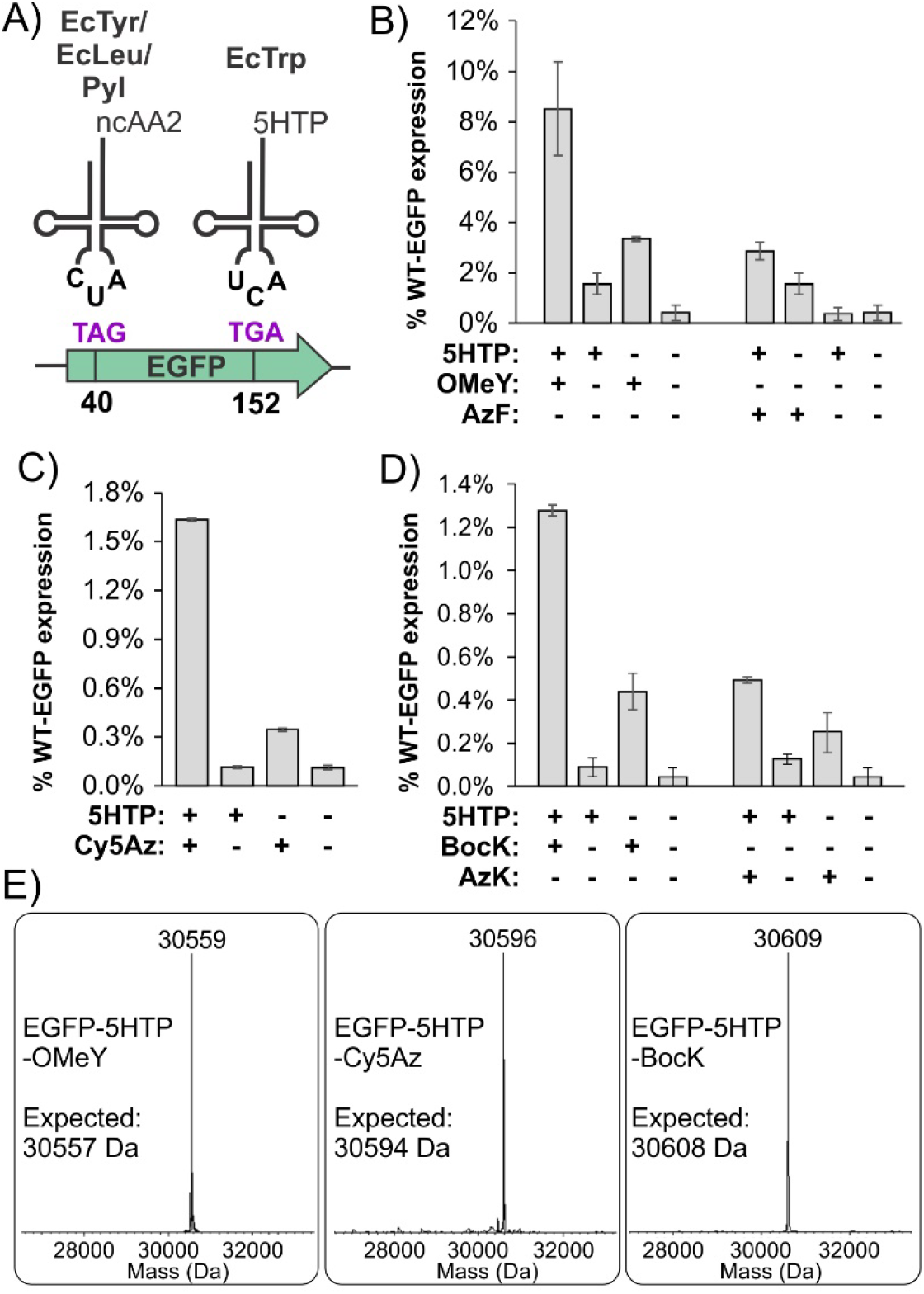
Three new dual suppression platforms in mammalian cells using the EcTrpRS/tRNA_UCA_^EcTrp^ pair. A) The dual suppression scheme. Expression of the EGFP-40-TAG-152-TGA reporter upon co-transfection of HEK293T cells with pAcBac3-EcTrp_TGA_-EGFP** + pAcBac1-EcTyr_TAG_ (panel B), pAcBac3-EcLeu_TAG_-EGFP** + pAcBac1-EcTrp_TGA_ (panel C), and pAcBac3-EcTrp_TGA_-EGFP** + pAcBac1-Pyl_TAG_ (panel D) and in the presence or absence of the indicated ncAAs. E) Deconvoluted ESI-MS analysis of purified EGFP reporters show masses consistent with the incorporation of the desired ncAAs.

Some of the ncAA combinations used above harbor two mutually compatible bioconjugation handles, which enable protein labeling at two distinct sites. To explore this possibility, we used the Pyl_TAG_+EcTrp_TGA_ system to express and purify the EGFP double mutant containing AzK and 5HTP. We have previously established that the azide and the 5-hydroxyindole groups in AzK and 5HTP can be labeled using strain-promoted azide alkyne cycloaddition (SPAAC) and the chemoselective rapid azo-coupling reaction (CRACR),^18,33–36^ respectively, without interfering with each other. Indeed, we demonstrated that the EGFP-AzK-5HTP expressed and purified from mammalian cells can be dually-labeled with two different fluorophores, DBCO-TAMRA and diazo-fluorescein (Figure S3), using SDS-PAGE (Figure 4B) and ESI-MS analysis (Figure 4C).

**Figure 4.**
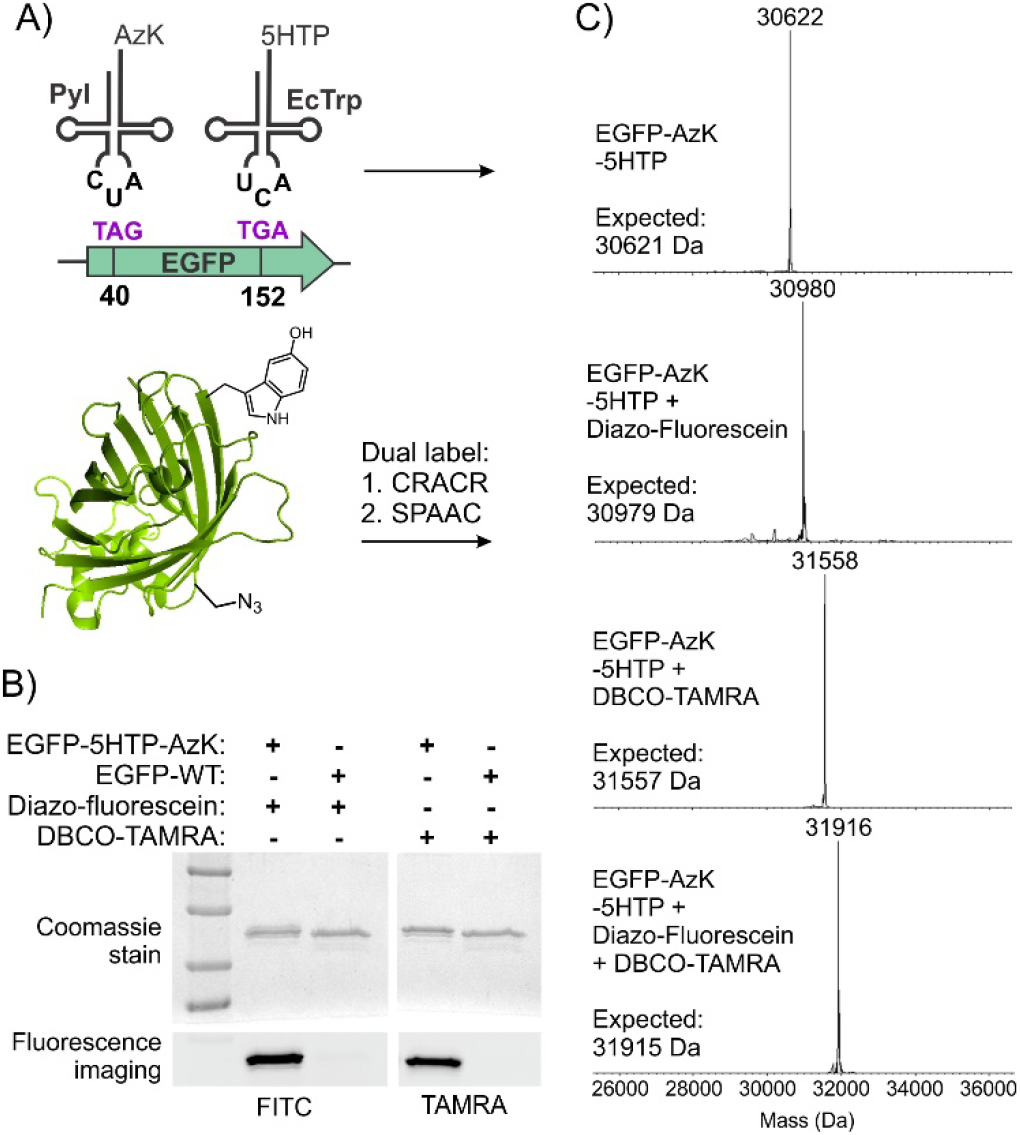
Site-specific dual labeling of EGFP-AzK-5HTP. A) The scheme of the experiment. B) Treatment with diazo-fluorescein or DBCO-TAMRA only results in the labeling of purified EGFP-AzK-5HTP, but not wild-type EGFP, as shown by fluorescent imaging following SDS-PAGE. C) MS analysis demonstrating efficient labeling of the 5HTP and AzK residues on EGFP-AzK-5HTP with diazo-fluorescein and DBCO-TAMRA, respectively, to achieve catalyst-free one-pot dual labeling.

Chemical modification of full-length antibodies has become a cornerstone for developing novel biotherapeutics, diagnostics, as well as research reagents.^7,37,38^ In this regard, the ability to introduce multiple distinct functionalities into antibodies with complete site-control can be transformative. Although it has been possible to use the EcTyr+Pyl dual suppression system to generate a full-length antibody containing two distinct ncAAs, the efficiency of expression was very low (100 μg/L).^27^ We surmised that the new dual suppression systems reported here, leveraging the efficient opal-suppressing EcTrp pair, can facilitate enhanced expression of antibodies site-specifically incorporating two distinct ncAAs. To this end, we attempted the expression of full-length Trastuzumab harboring a TAG and a TGA codon at positions 121 of the heavy chain (HC) and 169 of the light chain (LC), respectively, using the EcTrp+EcTyr system (Figure 5A), which showed the highest dual suppression efficiency. The antibody expression cassettes were incorporated into a vector containing a recently developed EcTyrRS (pAAFRS),^39^ capable of incorporating an azide-containing amino acid pAAF (**8**, Figure 1), as well as eight copies of tRNA_CUA_^EcTyr^. The pAAF incorporation system was selected due to its high efficiency, as well as the post-translational stability of this azide-ncAA within the reducing cellular environment.^39^ Expi293 suspension cultures were co-transfected with the resulting plasmid along with another plasmid encoding the EcTrpRS/tRNA_UCA_^EcTrp^ pair. A wild-type antibody, lacking any nonsense codons was also expressed in parallel. Full-length double mutant Trastuzumab-HC-121-pAAF-LC-169-5HTP was purified from the culture media by protein-G-affinity chromatography with impressive yields (6 mg/L; approximately 50% of wild-type Trastuzumab), a remarkable improvement over the previously reported system for dual-ncAA incorporation into an antibody (100 μg/L).^27^ ESI-MS analysis of the purified double mutant antibody confirmed light chain and heavy chain masses consistent with the incorporation of the desired ncAAs (Figure 5A). Treatment of this Trastuzumab double mutant with DBCO-TAMRA and diazo-fluorescein led to selective labeling of the HC and LC, respectively, while identical treatment of the wild-type protein led to no detectable labeling (Figure 5B), further confirming the presence of these unique bioconjugation handles at intended positions. The ability to efficiently express dual-ncAA labeled full-length antibodies will unlock many exciting applications, which we are actively exploring.

**Figure 5.**
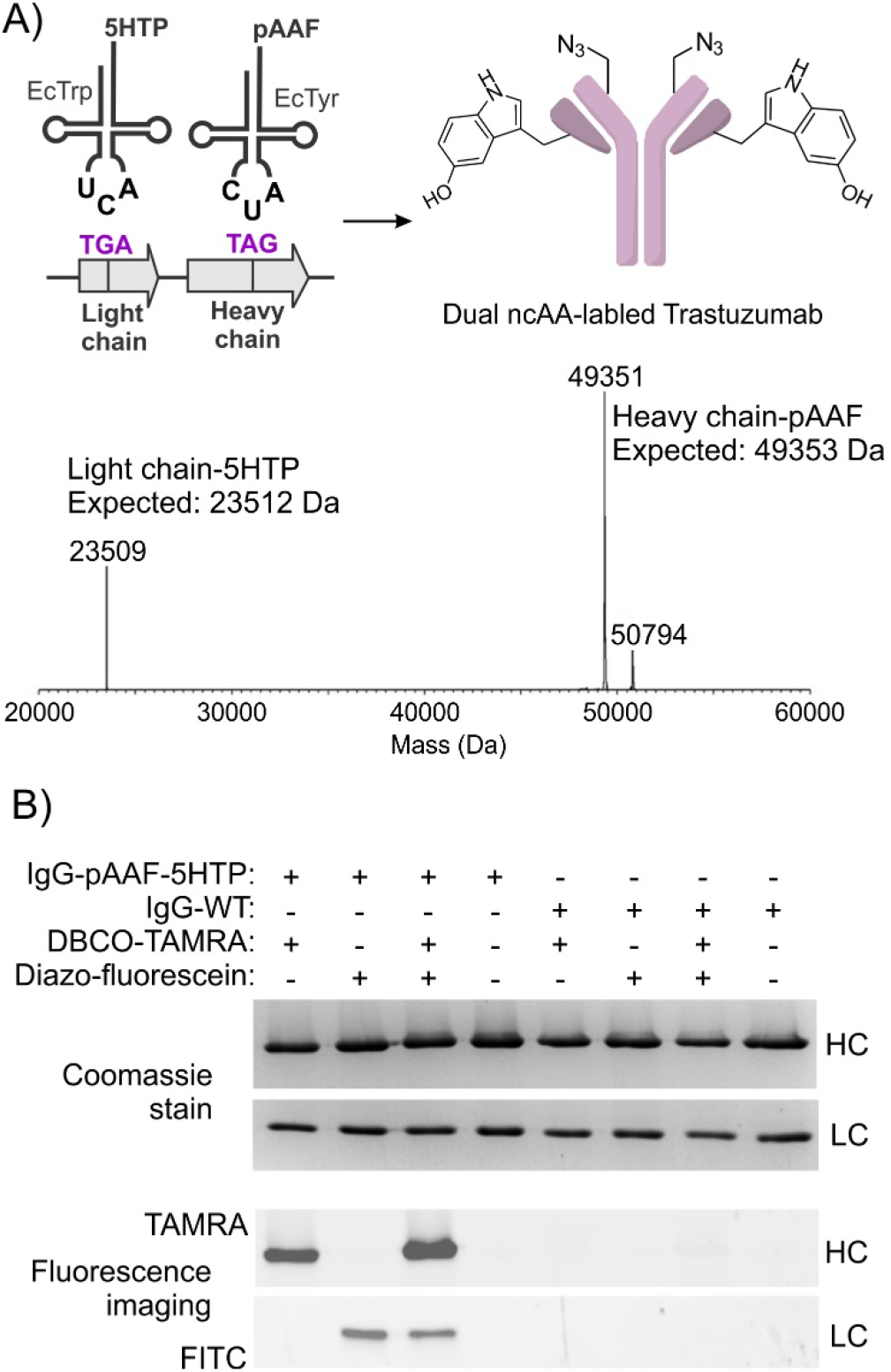
Incorporation of two distinct bioconjugation handles into full-length Trastuzumab using EcTrp+EcTyr dual suppression system. A) The scheme of the experiment. B) Deconvoluted ESI-MS analysis of full-length Trastuzumab (reduced, deglycosylated) expressed from Expi293 cells with 5HTP incorporated at position 169 of the light chain and pAAF incorporated at position 121 of the heavy chain using the EcTrp+EcTyr dual suppression system. The minor 50794 Da peak results from incomplete deglycosylation. C) Treatment with diazo-fluorescein or DBCO-TAMRA only results in the labeling of purified Trastuzumab-HC-pAAF-LC-5HTP, but not the wild-type antibody, as shown by fluorescent imaging following SDS-PAGE.

### Site-specific incorporation of three distinct ncAAs into one protein in mammalian cells

Demonstration that the EcTrp(TGA) pair does not cross-react with EcTyr and Pyl pairs, and given that EcTyr(TAG) and Pyl(TAA) pairs were previously established to be mutually orthogonal,^10,27^ it seemed plausible that these three pairs could be combined to facilitate site-specific incorporation of three distinct ncAAs into proteins in mammalian cells. The reassignment of all three nonsense codons does raise the issue of how to efficiently terminate translation. We have previously shown that three consecutive TAA codons can efficiently terminate translation in *E. coli*.^18^ To overcome any heterogeneity at the C-terminus from unwanted ncAA incorporation, we also developed a self-cleaving GTEV tag.^18^ In this system, the reporter protein is followed by a polyhistidine tag, a TEV cleavage site, TEV protease, and three UAA stop codons. Upon expression, the C-terminal TEV protease cleaves itself out, leaving the reporter protein with the purification tag and a clean C-terminus. To confirm that the GTEV expression system is functional in mammalian cells, we expressed wild-type EGFP from this system, and isolated it using the C-terminal polyhistidine tag. SDS-PAGE and ESI-MS analysis confirmed the production of the fully processed protein (Figure S4).

Next, we designed an expression system for incorporating three ncAAs simultaneously (Figure 6A), consisting of three plasmids containing: 1) A triple mutant EGFP reporter (EGFP-3-TAG-40-TGA-152-TAA) expressed from GTEV, and the EcTyrRS/tRNA_CUA_^EcTyr^ pair, 2) the EcTrpRS/tRNA_UCA_^EcTrp^ pair, and 3) the MbPylRS/tRNAU_AA_^Pyl^ pair. The third plasmid used an enhanced tRNA^Pyl^, recently developed in our group.^40^ These three plasmids were co-transfected into HEK293T cells and reporter expression was measured using fluorescence in the presence or absence of selected ncAAs. For EcTyr, either OMeYRS or pAAFRS were used for the incorporation of OMeY or para-amidoazidophenylalanine (pAAF, **8**; Figure 1), respectively. For Pyl, either BocK or cyclopropenelysine (CpK, **7**; Figure 1) were used, and 5HTP was used with the EcTrp system. Significant reporter expression was only observed when all three ncAAs were present for both OMeY+BocK+5HTP and pAAF+CpK+5HTP, indicating successful incorporation of the desired ncAAs (Figure 6B, Figure S8). The expression of the triple mutant proteins was approximately 1% of wild-type EGFP WT, expressed from the same GTEV vector. The fulllength triple mutant proteins were successfully isolated and analyzed by SDS-PAGE (Figure 6B) and whole-protein ESI-MS, which revealed masses consistent with the incorporation of the three desired ncAAs (Figure 6C-D).

**Figure 6.**
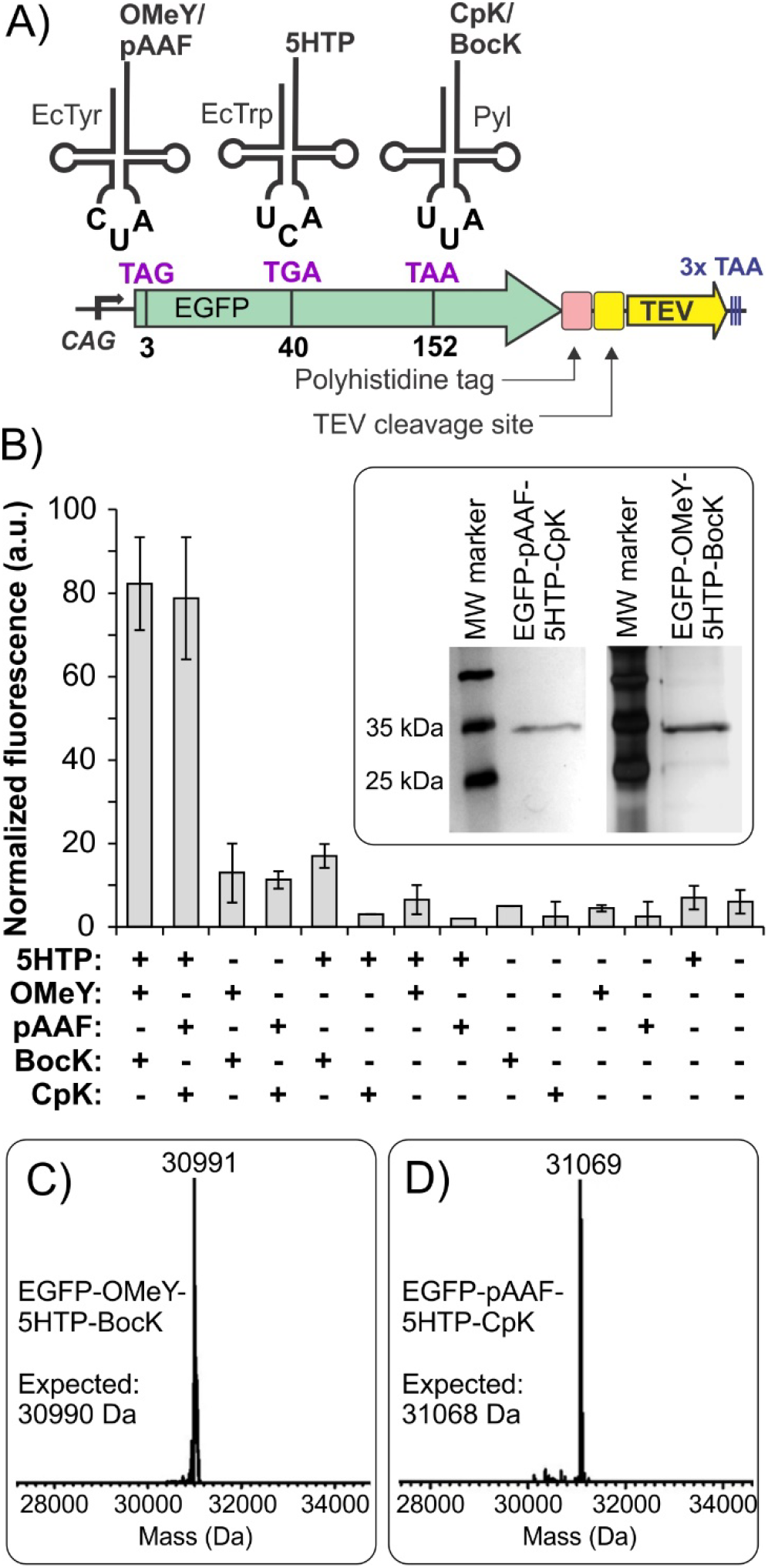
Site-specific incorporation of three distinct ncAAs into a protein in mammalian cells. A) The scheme describing the triple ncAA incorporation experiment that uses a TEV-based self-cleaving tag to generate a homogeneous C-terminus. B) Expression of EGFP-3TAG-40TGA-152TAA in the presence of the indicated ncAAs, measured by its characteristic fluorescence. Robust expression is observed only in the presence of three different ncAA substrates. Inset shows SDS-PAGE analyses of purified EGFP-BocK-OMeY-5HTP and EGFP-CpK-pAAF-5HTP. Deconvoluted whole protein MS analyses of EGFP-BocK-OMeY-5HTP (panel C) and EGFP-CpK-pAAF-5HTP (panel D) show masses consistent with the incorporation of the desired ncAAs.

In conclusion, here we show that the EcTrpRS/tRNA pair is an excellent opal suppressor in mammalian cells, and that it can be combined with amber-suppressing EcTyr, EcLeu, and Pyl pairs without cross-reaction to create three new ways to site-specifically incorporate two distinct ncAAs into proteins. Each of these platforms are already populated with many useful ncAAs, and more can be readily introduced through facile directed evolution systems. The EcTrp+EcTyr platform enabled remarkably efficient incorporation of two distinct ncAAs – with mutually compatible bioconjugation handles – into a full-length humanized antibody. Finally, we showed that the EcTrp, EcTyr, and Pyl platforms can be combined to site-specifically incorporate three distinct ncAAs into proteins expressed in mammalian cells. Although the yield of the triply-labeled protein was low, the efficiency of this first-generation technology can be improved through expression plasmid optimization, as well as enhancing the efficiency of the aaRS/tRNA pairs.^39,40^ The work described here significantly expands the scope of multi-ncAA mutagenesis in mammalian systems and would facilitate numerous applications.

## Acknowledgements

This work was supported by NIH grant R35GM136437 to A.C.

## Additional information

Supplementary information is available for this paper.

