## Supporting information for "An efficient opal-suppressor tryptophanyl pair creates new routes for simultaneously incorporating up to three distinct noncanonical amino acids into proteins in mammalian cells"

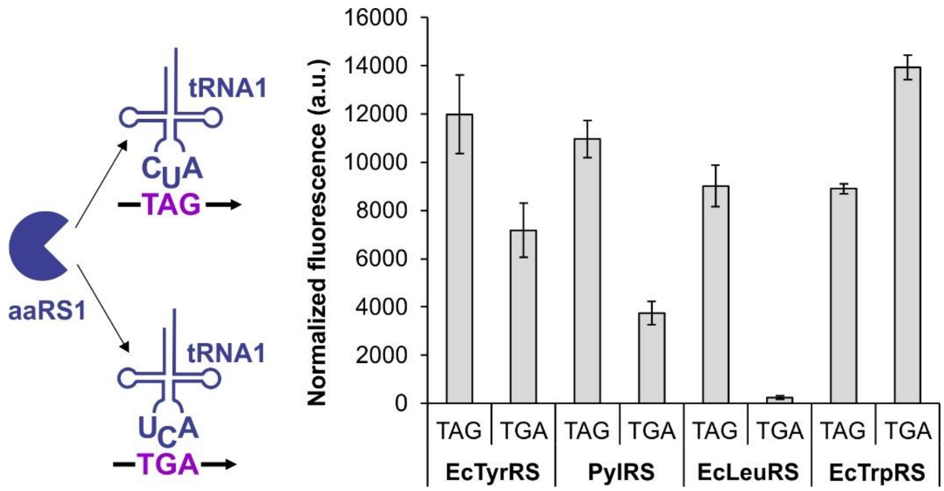


**Figure S1**. EcTrpRS/tRNA pair is an excellent opal suppressor. Each pair was tested for opal and amber suppression efficiency in HEK293T cells with an EGFP reporter harboring an appropriate nonsense codon at the permissive position 40.

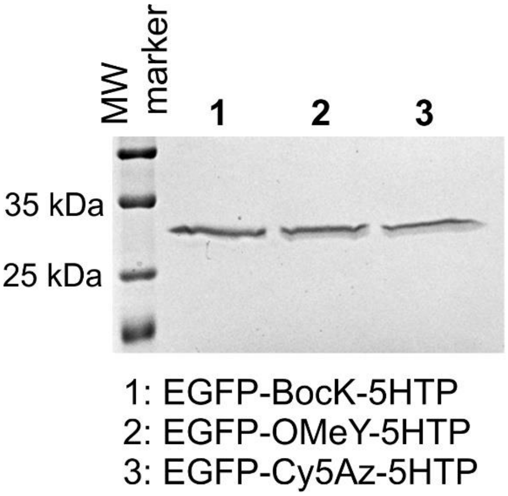


**Figure S2**. SDS-PAGE analysis of purified EGFP reporter proteins from the dual suppression experiments described in Figure 3.

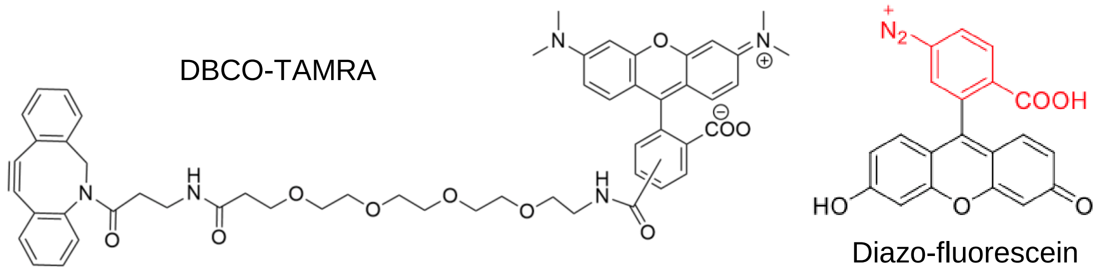


**Figure S3**. Structures of DBCO-TAMRA and diazo-fluorescein.

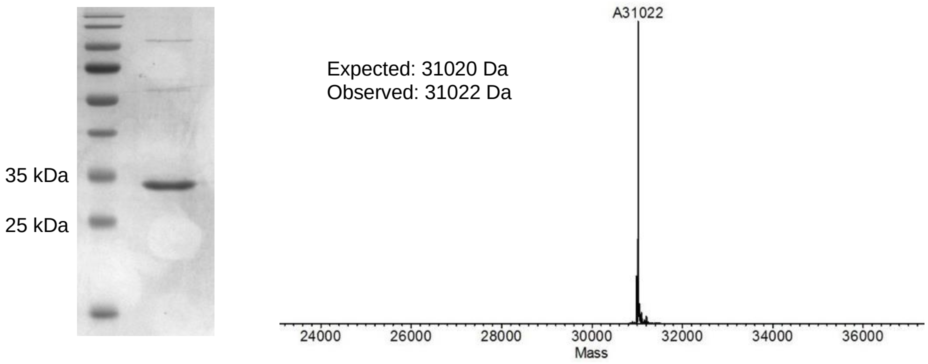


**Figure S4.** Purification of wild-type EGFP, expressed using the GTEV system in HEK293T cells. Cells were transfected with pAcBac3-EGFP-WT-TEV-3xUAA, harvested 48 h post-transfection, and EGFP was purified using the C-terminal polyhistidine tag. The purified protein was subjected to SDS-PAGE analysis and ESI-MS to show complete removal of the C-terminal self-cleaving tag.

### Materials and Methods

#### General methods:

All cloning and plasmid propagation were performed in *E. coli* strain DH10b. Antibiotics were supplied in the bacterial culture media at the following concentrations: 150 μg/mL ampicillin, 50 μg/mL kanamycin. Oligonucleotides were purchased from Azenta Life Sciences. Phusion polymerase, restriction enzymes, and T4 DNA ligase were obtained from Fisher Scientific. Sanger sequencing of DNA was performed by Azenta Life Sciences. For mammalian cell culture, HEK293T cells were maintained at 37 °C and 5% $CO_2$ using DMEM supplemented with 10% fetal bovine serum and penicillin-streptomycin at a final concentration of 100 U/mL. High glucose DMEM, trypsin 0.25%, penicillin-streptomycin (10,000 U/mL), and fetal bovine serum (FBS) were obtained from Fisher Scientific. Polyethylenimine PEI MAX from Polysciences (Warrington, PA) was used for transient transfection. Sodium butyrate used for transfection was obtained from Fisher Scientific.

O-methyl-l-tyrosine (OMeY) and $N^{\varepsilon}$-boc-l-lysine (BocK) were obtained from Fisher Scientific. 5-hydroxy-l-tryptophan (5HTP) and 4-azido-l-phenylalanine (AzF) were obtained from Chem Impex. Cyclopropene-l-lysine (CpK) was obtained from Sirius Fine Chemicals. $N^{\varepsilon}$-azido-l-lysine (AzK) was obtained from Iris Biotech. Cy5Az and pAAF were synthesized as described previously.[2,6]

#### Plasmid construction for expressing aaRS, tRNA, and EGFP* separately:

Plasmids for expressing each aaRS, tRNA, and EGFP mutants individually were previously reported for the EcTyr, Pyl, and EcLeu systems.[1-2] Plasmid for expressing the evolved *E. coli* tryptophanyl synthetase, pb1-EcWRS-h14 was also previously reported.[3] The TAG suppressing tryptophanyl tRNA was cloned into a pIDTSmart vector following a previously described protocol using NheI/AvrII restriction digest.[3] Overlap extension PCR was performed on the subsequent plasmid to produce the equivalent TGA suppressing tRNA using primers pIDTSmart-F, Cargo-R, EcWtR-TGA-iF, EcWtR-TGA-iR, and NheI/AvrII restriction enzyme digest.

#### Plasmids for dual suppression:

To create pAcBac1-$EcTrp_{TGA}$, first a cassette containing 8 copies of the TGA suppressing EcWtR was constructed following a previously reported protocol.[4] The tRNA cassette was then digested with NheI/AvrII and inserted into the SpeI site of the previously reported pb1-EcWRS-h14 plasmid.[3] To construct pAcBac3-$EcTrp_{TGA}$-EGFP**, a 4 copy cassette of TGA suppressing EcWtR was constructed and inserted into the pb1-EcWRS-h14 plasmid, and EGFP-40TGA-152TAG was inserted using SfiI restriction enzyme digest as described previously.[1] pAcBac3-$EcLeu_{TAG}$-EGFP**, pAcBac1-$EcTyr_{TAG}$ , and pAcBac1-$Pyl_{TAG}$ were reported previously.[1,2]

For expression of Trastuzumab, pcDNA3.1-Her2 plasmid (gift from Prof. Han Xiao) was first mutagenized using primer HugI-121A-TAG and PrimeSTAR MAX DNA polymerase (Takara) to clone a TAG codon into position 121 of the heavy chain. The subsequent plasmid was further mutagenized using primer LC-169-TGA-F and PrimeSTAR MAX DNA polymerase, adding a TGA codon at position 169 of the light chain, to create pcDNA3.1-Her2-HC121TAG-LC169TGA. The antibody expression cassette was then amplified out of that plasmid using anti-Her2-BamHI-F and anti-Her2-SfiI-R primers, digested with BamHI and SfiI, and inserted into a pacbac3 plasmid containing 8 copies of $tRNA_{CUA}^{EcTyr}$ and pAAFRS[6], replacing the EGFP expression cassette.

**Plasmids for triple suppression:**

Site-directed mutagenesis was used to construct pb1-EGFP-3TAA-40TGA-152TAG by first amplifying pb1-EGFP-40TGA with mutagenic primer EGFP-152-TAG-F and PrimeSTAR Max DNA polymerase (Takara) to create pb1-EGFP-40TGA-152TAG, which was subsequently amplified using mutagenic primer EGFP-3TAA-F and PrimeSTAR Max DNA polymerase to create the triple mutant.

To create pacbac3-OMeYRS-8xYtR-EGFP-3TAA-40TGA-152TAG-TEV-3xUAA, an overlap extension PCR was performed, stitching together the EGFP with polyhistidine tag from pb1-EGFP-3TAA-40TGA-152TAG, and the TEV cleavage site and TEV protease followed by 3xUAA from previously reported GTEV plasmid[5] using primers EGFP-3TAA-SfiI-F, EGFP-10xHis-TEVoverlap-R, TEV-overlap-F, TEV-3xUAA-SfiI-R. The PCR product was digested with SfiI and inserted into previously reported pAcBac3 plasmid containing OMeYRS and 8xEcYtR, replacing the EGFP. To create the analogous pAcBac3-pAAFRS-8xYtR-EGFP-3TAA-40TGA-152TAG-TEV-3xUAA plasmid, a previously reported plasmid containing EcTyrRS mutant pAAFRS[6] and pAcBac3-OMeYRS-8xYtR-EGFP-3TAA-40TGA-152TAG-TEV-3xUAA were both digested with NheI/XhoI followed by ligation.

The analogous wild-type GTEV plasmid, pAcBac3-EGFP-WT-TEV-3xUAA, was constructed using the same overlap extension method but using primer EGFP-SfiI-F and amplifying EGFP from pb1-EGFP-WT.

**Analysis of EGFP reporter expression in HEK293T cells:**

For small-scale EGFP expression, HEK293T cells were seeded at a density of 600,000 cells/ well in a 12-well plate one day prior to transfection. A total amount of 1 μg DNA + 4 μL PEI (1 mg/mL) + 20 μL DMEM was used for transfection of each well. For two-plasmid transfection, 0.5 ug of each plasmid was used. For three-plasmid transfection, a total of 1.2 μg of DNA was used, 0.4 μg of each plasmid. The appropriate ncAAs were supplied to each well at a final concentration of 1 mM. Fluorescence analysis was performed 48 hrs post-transfection. To obtain fluorescence data, cells were harvested and lysed as previously described.[2] Fluorescence data was collected in a 96-well plate using a SpectraMAX M5 (Molecular Devices) (ex = 488 nm and em = 510 nm). Mean of three independent experiments were reported, and error bars represent standard deviation. Fluorescence images were taken on a Zeiss Axio Observer fluorescence microscope.

**Expression and purification of EGFP double mutants:**

For larger scale protein expression incorporating two ncAAs, HEK293T cells were seeded in 100 mm cell culture dishes (8 million cells per dish) one day prior to transfection. A total amount of 12 μg DNA (6 μg of each plasmid) + 48 μl PEI (1 mg/mL) + 240 μl DMEM was used to transfect each dish. The ncAAs were supplemented at 1 mM final concentration at the time of transfection. Sodium butyrate was also supplied in the transfection media at a final concentration of 2 mM. Cells were harvested 48 h post-transfection, lysed with CelLytic M, and protein was purified from the clarified lysate via the C-terminal polyhistidine tag using HisPur Ni-NTA resin following the manufacturer's protocol. Purified proteins were subjected to SDS-PAGE analysis and ESI-MS (Agilent TOF HPLC-MS).

**EGFP double mutant protein labeling:**

To dually label EGFP-AzK-5HTP, a previously described protocol was followed.[5] Briefly, 10 μM EGFP-AzK-5HTP was first labeled with 50 μM diazo-fluorescein for 30 min at RT and quenched with 1 mM free 5HTP. Next, the protein was further labeled in the same pot using 200 μM DBCO-TAMRA (Click Chemistry Tools) for 3 h at ambient temperature and quenched afterward with 1 mM free AzK. The resulting protein was then subjected to SDS-PAGE followed by fluorescence imaging (ChemiDoc MP, Bio-Rad), as well as ESI-MS (Agilent TOF HPLC-MS) analysis. Diazo-fluorescein was synthesized as described previously.[7]

**Antibody expression and purification:**

Expi293F cells were grown at 37 °C, 8% $CO_2$, on an orbital shaker at 125 rpm in Expi293 expression media supplemented with 0.5x antibiotic-antimycotic (Fisher). Cells were seeded at a density of 0.3 million cells/mL and sub-cultured every four days. For transfection, when cells reached a density of 3 million cells/mL (at least 95% viability), they were spun down and resuspended in media to a density of 20 million cells/mL. Equal ratio of the two plasmids were added to this cell suspension at a final DNA concentration of 25 μg/mL. PEI MAX (40 mg/mL stock) was added to the cells at a final concentration of 50 μg/mL. After addition of DNA and PEI, cells were incubated under standard conditions with shaking for 3 hours, then diluted back to their original volume. The appropriate ncAAs were each supplied at a final concentration of 1 mM, and 2 mM valproic acid was also added.

After 7-10 days of expression, the cells were pelleted, and the decanted media was passed through a 0.22 μm sterile filter. The solution was adjusted to pH 5.4, 50 mM NaOAc. After 1 mL of Pierce™ Protein G Agarose resin (Thermo Scientific) was equilibrated with wash buffer (50 mM NaOAc pH 5.4), the media was passed through by gravity flow. The column was then washed with 50 mM NaOAc pH 5.4, with a volume equal to approximately half the volume of media passed through the column. The antibody was then eluted by 12 stepwise additions of 1 mL of 100 mM glycine (pH 2.7) into tubes containing phosphate buffer pH 8 to neutralize the fractions immediately. The elution fractions were concentrated using an Amicon Ultra 15 centrifugal filter unit and exchanged to PBS buffer. Protein concentration was determined by $A_{280}$ using NanoDrop 2000 spectrophotometer. For ESI-MS analysis, the antibody was first reduced with 10 mM TCEP at 55 °C for 15 min, followed by treatment with Remove-iT PNGaseF (New England BioLabs) for 1 h at 37 °C. The PNGase was removed by incubation with chitin resin (New England BioLabs) for 30 min at RT, centrifuged to remove the resin, and the supernatant was subjected to ESI-MS (Agilent TOF HPLC-MS) analysis.

**EGFP triple mutant expression and purification:**

HEK293T cells were seeded in 100 mm cell culture dishes (8 million cells per dish) one day prior to transfection. A total amount of 12 μg DNA (4 μg of each plasmid) + 48 μL PEI (1 mg/mL) + 240 μL DMEM was used to transfect each dish. The three ncAAs were supplied in the media at the time of transfection, each at a final concentration of 1 mM. Sodium butyrate was also supplied at the time of transfection, at a final concentration of 2 mM. 48 h post-transfection, cells were harvested, lysed with CelLytic M, and the triple mutant EGFP was purified from the clarified lysate using GFP-Trap Agarose (Chromotek) resin following the manufacturers protocol. Following SDS-PAGE, the proteins were visualized by silver staining using ProteoSilver Silver Staining Kit (Millipore Sigma) and subjected to ESI-MS (Agilent TOF HPLC-MS).

**Table S1.** Oligonucleotide sequences.

| Name | Sequence |
|---|---|
| EGFP-SfiI-F | AATAATGAATTGGCCAAGGAGGCCACCATGGTG |
| EGFP-3TAA-SfiI-F | aatatggccaaggaggccGCCGCCAcCATGTAAGTGAG |
| EGFP-10xHis-TEVoverlap-R | ttgaaaataaagattttcgtgatggtgatgATGGTGATGGTGATGATGACCGGTATGC |
| TEV-overlap-F | caccatcatcaccatcacGAAAATCTTTATTTTCAAggtggaggaagtggagaaag |
| TEV-3xUAA-SfiI-R | taataggccttagaggccTTATTAttagcgacggcgacgacgattcatgag |
| EGFP-3TAA-F | AATAATGCCGCCAcCATGGTGTAAAGCAAGGGCGAGGAGC |
| EGFP-152TAG-F | AATAATCAACAGCCACAACGTCTAGATCATGGCCGACAAGCAGAAG |
| EcWtR-TGA-iF | CAaCACCCGGTTTTgaAGACCGGTGCTCTACCAATTGAACTAC |
| EcWtR-TGA-iR | CCGGTCTTcAAAACCGGGTGtTGGGAGTTCG |
| pIDTSmart-F | GTTGCGTTTGAGACGGGCGA |
| Cargo-R | GTCCAGTAGTGATCGACACTGCTCG |
| anti-Her2-BamHI-F | AATAATGGATCCcgcggtgagttcaggc |
| anti-Her2-SfiI-R | TTATTAGGCCTTAGAGGCCGCttaAGAacccg |
| HugI-121A-TAG | gggcaccctggtcacagtgtcctctTAGagcaccaagggcccatcggtc |
| LC-169-TGA-F | aataatgtcacagagcaggacagctgagacagcacctacagcctc |

**Plasmid sequences:**

pIDTSmart-1xU6-EcWtR-TGA: EcWtR is colored blue, and U6 promoter is colored purple.

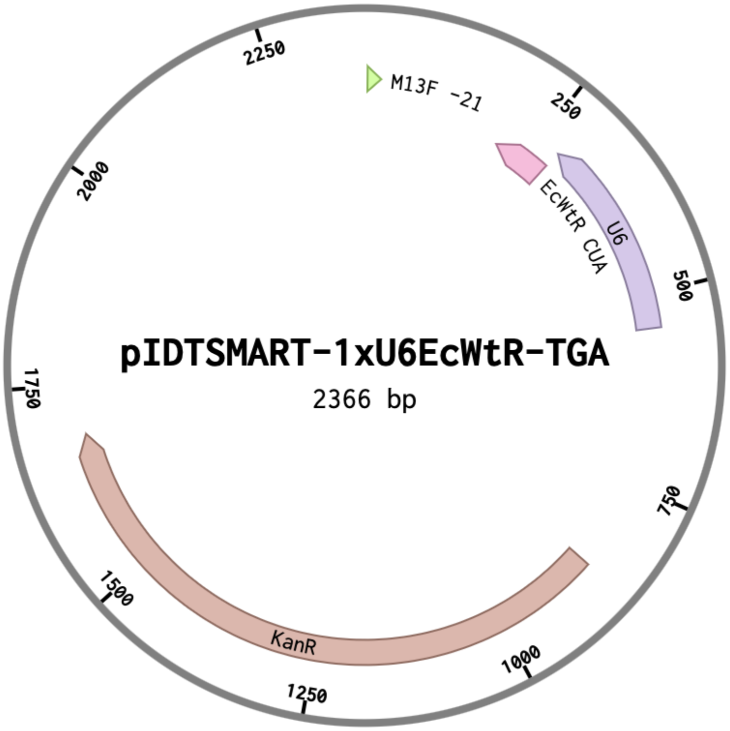


CCCGTGTAAAACGACGGCCAGTTTATCTAGTCAGCTTGATTCTAGCTGATCGTGGAC
CGGAAGGTGAGCCAGTGAGTTGATTGCAGTCCAGTTACGCTGGAGTCTGAGGCTCGT
CCTGAATGATATGCGaCCGCCGGAGGGTTGCGTTTGAGACGGGCGACAGATCCAGTC
GCGCTGCTCTCGTCGATCCGctagcaaaaaaTGGCAGGGGCGGAGAGACTCGAACTCCCAa
CACCCGGTTTTgaAGACCGGTGCTCTACCAATTGAACTACGCCCCTGGTGTTTCGTCC
TTTCCACAAGATATATAAAGCCAAGAAATCGAAATACTTTCAAGTTACGGTAAGCAT
ATGATAGTCCATTTTAAAACATAATTTTAAAACTGCAAACTACCCAAGAAATTATTA
CTTTCTACGTCACGTATTTTGTACTAATATCTTTGTGTTTACAGTCAAATTAATTCTA
ATTATCTCTCTAACAGCCTTGTATCGTATATGCAAATATGAAGGAATCATGGGAAAT
AGGCCCTCTTCCTGCCCGACCTAGGGGTGCGAGCGGATCGAGCAGTGTCGATCACTA
CTGGACCGCGAGCTGTGCTGCGACcCGTGATCTTACGGCATTATACGTATGATCGGT
CCACGATCAGCTAGATTATCTAGTCAGCTTGATGTCATAGCTGTTTCCTGAGGCTCA
ATACTGACCATTTAAATCATACCTGACCTCCATAGCAGAAAGTCAAAAGCCTCCGAC
CGGAGGCTTTTGACTTGATCGGCACGTAAGAGGTTCCAACTTTCACCATAATGAAAT
AAGATCACTACCGGGCGTATTTTTTGAGTTATCGAGATTTTCAGGAGCTAAGGAAGC
TAAAATGAGCCATATTCAACGGGAAACGTCTTGCTTGAAGCCGCGATTAAATTCCAA
CATGGATGCTGATTTATATGGGTATAAATGGGCTCGCGATAATGTCGGGCAATCAGG
TGCGACAATCTATCGATTGTATGGGAAGCCCGATGCGCCAGAGTTGTTTCTGAAACA
TGGCAAAGGTAGCGTTGCCAATGATGTTACAGATGAGATGGTCAGGCTAAACTGGC
TGACGGAATTTATGCCTCTTCCGACCATCAAGCATTTTATCCGTACTCCTGATGATGC
ATGGTTACTCACCACTGCGATCCCAGGGAAAACAGCATTCCAGGTATTAGAAGAAT

ATCCTGATTCAGGTGAAAATATTGTTGATGCGCTGGCAGTGTTCCTGCGCCGGTTGC
ATTCGATTCCTGTTTGTAATTGTCCTTTTAACGGCGATCGCGTATTTCGTCTCGCTCA
GGCGCAATCACGAATGAATAACGGTTTGGTTGGTGCGAGTGATTTTGATGACGAGC
GTAATGGCTGGCCTGTTGAACAAGTCTGGAAAGAAATGCATAAACTCTTGCCATTCT
CACCGGATTCAGTCGTCACTCATGGTGATTTCTCACTTGATAACCTTATTTTTGACGA
GGGGAAATTAATAGGTTGTATTGATGTTGGACGAGTCGGAATCGCAGACCGATACC
AGGATCTTGCCATCCTATGGAACTGCCTCGGTGAGTTTTCTCCTTCATTACAGAAAC
GGCTTTTTCAAAAATATGGTATTGATAATCCTGATATGAATAAATTGCAGTTTCACTT
GATGCTCGATGAGTTTTTCTAATGAGGACCTAAATGTAATCACCTGGCTCACCTTCG
GGTGGGCCTTTCTGCGTTGCTGGCGTTTTTCCATAGGCTCCGCCCCCCTGACGAGCAT
CACAAAAATCGATGCTCAAGTCAGAGGTGGCGAAACCCGACAGGACTATAAAGATA
CCAGGCGTTTCCCCCTGGAAGCTCCCTCGTGCGCTCTCCTGTTCCGACCCTGCCGCTT
ACCGGATACCTGTCCGCCTTTCTCCCTTCGGGAAGCGTGGCGCTTTCTCATAGCTCAC
GCTGTAGGTATCTCAGTTCGGTGTAGGTCGTTCGCTCCAAGCTGGGCTGTGTGCACG
AACCCCCCGTTCAGCCCGACCGCTGCGCCTTATCCGGTAACTATCGTCTTGAGTCCA
ACCCGGTAAGACACGACTTATCGCCACTGGCAGCAGCCACTGGTAACAGGATTAGC
AGAGCGAGGTATGTAGGCGGTGCTACAGAGTTCTTGAAGTGGTGGCCTAACTACGG
CTACACTAGAAGAACAGTATTTGGTATCTGCGCTCTGCTGAAGCCAGTTACCTCGGA
AAAAGAGTTGGTAGCTCTTGATCCGGCAAACAAACCACCGCTGGTAGCGGTGGTTTT
TTTGTTTGCAAGCAGCAGATTACGCGCAGAAAAAAAGGATCTCAAGAAGATCCTTT
GATTTTCTACCGAAGAAAGGCCCA

**pb3-EcWRS-h14-4xU6-EcWtR-TGA-EGFP**:** EcWRS-h14 is colored in magenta, EcWtR is colored in blue, U6 promoter is colored in purple, and EGFP is colored green.

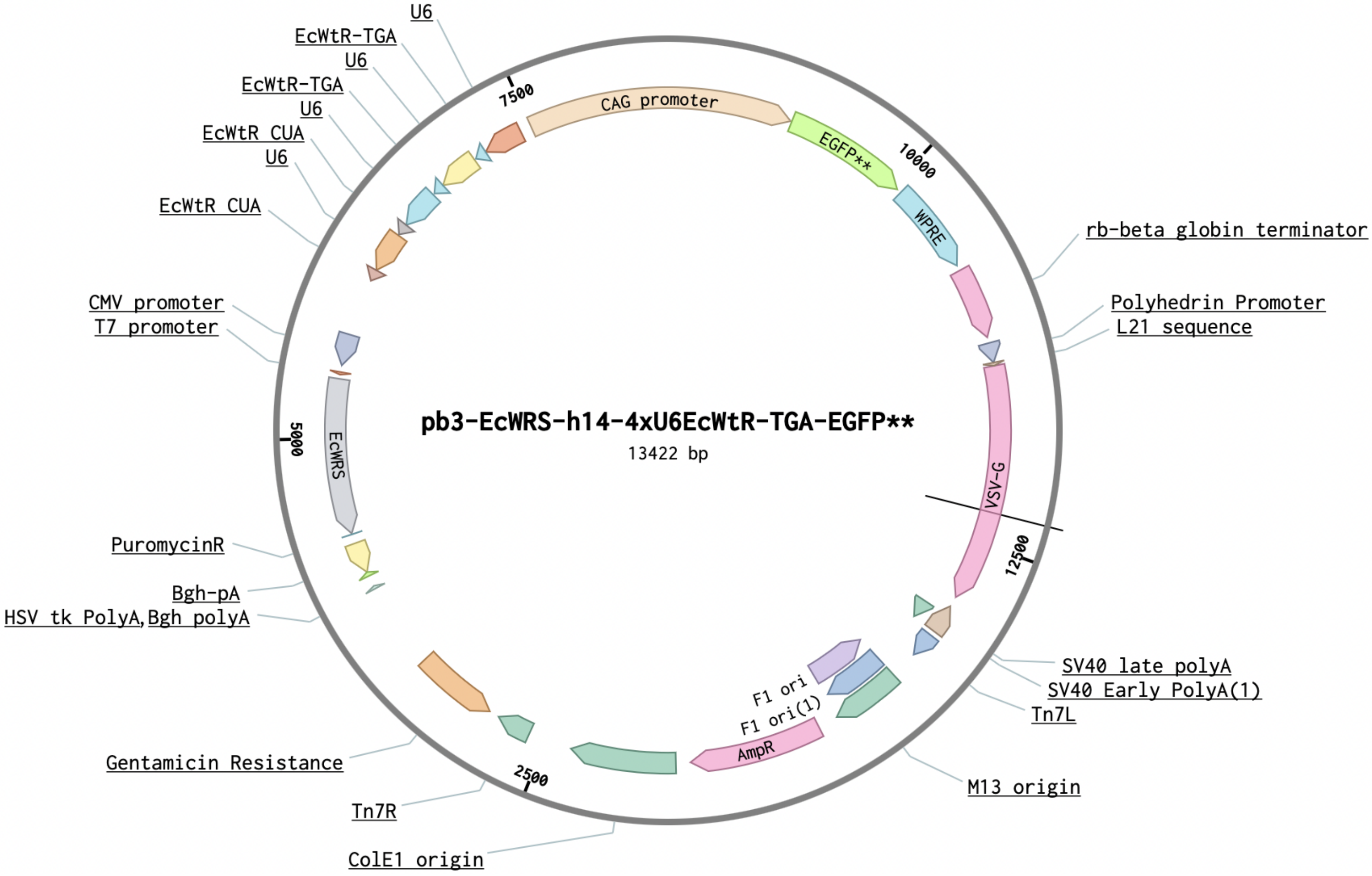


ttctctgtcacagaatgaaaatttttctgtcatctcttcgttattaatgtttgtaattgactgaatatcaacgcttatttgcagcctgaatggcgaatgg
gacgcgccctgtagcggcgcattaagcgcggcgggtgtggtggttacgcgcagcgtgaccgctacacttgccagcgccctagcgcccgc
tcctttcgctttcttcccttcctttctcgccacgttcgccggctttccccgtcaagctctaaatcgggggctccctttagggttccgatttagtgcttt
acggcacctcgaccccaaaaaacttgattagggtgatggttcacgtagtgggccatcgccctgatagacggtttttcgccctttgacgttgga
gtccacgttctttaatagtggactcttgttccaaactggaacaacactcaaccctatctcggtctattcttttgatttataagggattttgccgatttc
ggcctattggttaaaaaatgagctgatttaacaaaaatttaacgcgaattttaacaaaatattaacgtttacaatttcaggtggcacttttcgggga
aatgtgcgcggaacccctatttgtttatttttctaaatacattcaaatatgtatccgctcatgagacaataaccctgataaatgcttcaataatattg
aaaaaggaagagtatgagtattcaacatttccgtgtcgcccttattcccttttttgcggcattttgccttcctgtttttgctcacccagaaacgctgg
tgaaagtaaaagatgctgaagatcagttgggtgcacgagtgggttacatcgaactggatctcaacagcggtaagatccttgagagttttcgcc
ccgaagaacgttttccaatgatgagcacttttaaagttctgctatgtggcgcggtattatcccgtattgacgccgggcaagagcaactcggtcg
ccgcatacactattctcagaatgacttggttgagtactcaccagtcacagaaaagcatcttacggatggcatgacagtaagagaattatgcag
tgctgccataaccatgagtgataacactgcggccaacttacttctgacaacgatcggaggaccgaaggagctaaccgcttttttgcacaacat
gggggatcatgtaactcgccttgatcgttgggaaccggagctgaatgaagccataccaaacgacgagcgtgacaccacgatgcctgtagc
aatggcaacaacgttgcgcaaactattaactggcgaactacttactctagcttcccggcaacaattaatagactggatggaggcggataaagt
tgcaggaccacttctgcgctcggcccttccggctggctggtttattgctgataaatctggagccggtgagcgtgggtctcgcggtatcattgc
agcactggggccagatggtaagccctcccgtatcgtagttatctacacgacggggagtcaggcaactatggatgaacgaaatagacagat

cgctgagataggtgcctcactgattaagcattggtaactgtcagaccaagtttactcatatatactttagattgatttaaaacttcatttttaatttaa
aaggatctaggtgaagatcctttttgataatctcatgaccaaaatcccttaacgtgagttttcgttccactgagcgtcagaccccgtagaaaaga
tcaaaggatcttcttgagatcctttttttctgcgcgtaatctgctgcttgcaaacaaaaaaaccaccgctaccagcggtggtttgtttgccggatc
aagagctaccaactctttttccgaaggtaactggcttcagcagagcgcagataccaaatactgtccttctagtgtagccgtagttaggccacc
acttcaagaactctgtagcaccgcctacatacctcgctctgctaatcctgttaccagtggctgctgccagtggcgataagtcgtgtcttaccgg
gttggactcaagacgatagttaccggataaggcgcagcggtcgggctgaacggggggttcgtgcacacagcccagcttggagcgaacg
acctacaccgaactgagatacctacagcgtgagcattgagaaagcgccacgcttcccgaagggagaaaggcggacaggtatccggtaag
cggcagggtcggaacaggagagcgcacgagggagcttccagggggaaacgcctggtatctttatagtcctgtcgggtttcgccacctctg
acttgagcgtcgatttttgtgatgctcgtcaggggggcggagcctatggaaaaacgccagcaacgcggcctttttacggttcctggccttttg
ctggccttttgctcacatgttctttcctgcgttatcccctgattctgtggataaccgtattaccgcctttgagtgagctgataccgctcgccgcagc
cgaacgaccgagcgcagcgagtcagtgagcgaggaagcggaagagcgcctgatgcggtattttctccttacgcatctgtgcggtatttcac
accgcagaccagccgcgtaacctggcaaaatcggttacggttgagtaataaatggatgccctgcgtaagcgggtgtgggcggacaataaa
gtcttaaactgaacaaaatagatctaaactatgacaataaagtcttaaactagacagaatagttgtaaactgaaatcagtccagttatgctgtga
aaaagcatactggacttttgttatggctaaagcaaactcttcattttctgaagtgcaaattgcccgtcgtattaaagaggggcgtggccaaggg
catggtaaagactatattcgcggcgttgtgacaatttaccgaacaactccgcggccgggaagccgatctcggcttgaacgaattgttaggtg
gcggtacttgggtcgatatcaaagtgcatcacttcttcccgtatgcccaactttgtatagagagccactgcgggatcgtcaccgtaatctgcttg
cacgtagatcacataagcaccaagcgcgttggcctcatgcttgaggagattgatgagcgcggtggcaatgccctgcctccggtgctcgccg
gagactgcgagatcatagatatagatctcactacgcggctgctcaaacctgggcagaacgtaagccgcgagagcgccaacaaccgcttct
tggtcgaaggcagcaagcgcgatgaatgtcttactacggagcaagttcccgaggtaatcggagtccggctgatgttgggagtaggtggcta
cgtctccgaactcacgaccgaaaagatcaagagcagcccgcatggatttgacttggtcagggccgagcctacatgtgcgaatgatgcccat
acttgagccacctaactttgttttagggcgactgccctgctgcgtaacatcgttgctgctgcgtaacatcgttgctgctccataacatcaaacat
cgacccacggcgtaacgcgcttgctgcttggatgcccgaggcatagactgtacaaaaaaacagtcataacaagccatgaaaaccgccact
gcgccgttaccaccgctgcgttcggtcaaggttctggaccagttgcgtgagcgcatacgctacttgcattacagtttacgaaccgaacaggc
ttatgtcaactgggttcgtgccttcatccgtttccacggtgtgcgtcacccggcaaccttgggcagcagcgaagtcgaggcatttctgtcctgg
ctggcgaacgagcgcaaggtttcggtctccacgcatcgtcaggcattggcggccttgctgttcttctacggcaaggtgctgtgcacggatct
gccctggcttcaggagatcggtagacctcggccgtcgcggcgcttgccggtggtgctgaccccggatgaagtggttcgcatcctcggttttc
tggaaggcgagcatcgtttgttcgcccaggactctagctatagttctagtggttggcctacgtacccgtagtggctatggcagggcttgcgctt
aatgcgccgctacagggcgcgtggggataccccctagagccccagctggttctttccgcctcagaagccatagagcccaccgcatcccca
gcatgCCTGCTATTGTCTTCCCAATCCTCCCCCTTGCTGTCCTGCCCCACCCCACCCCCC
AGAATAGAATGACACCTACTCAGACAATGCGATGCAATTTCCTCATTTTATTAGGAA
AGGACAGTGGGAGTGGCACCTTCCAGGGTCAAGGAAGGCACGGGGGAGGGGCAAA
CAACAGATGGCTGGCAACTAGAAGGCACAGTCGAGGCTGATCAGCGGGTTTAAACG
GGCCCTCTAGACtcgagttaaagtcgacttaacgcgttgaattcTTACGGCTTCGCCACAAAACCAATCG
CTTCGTACACCGCTTTTAGCGTACGGGAAGCGTGCGCGCTGGCTTTTTCCGCGCCAT
CTTTCATCACCTGTTGCAGGAAGGCTTCATCGTTGCGGAAACGGTGATAGCGTTCCT
GCAATTCAGTCAGCATACCGGAAACGGCATCAGCCACTTCACCTTTCAGATGACCAT
ACATCTTGCCTTCGAACTGTTTTTCCAGTTCTGGGATGCTCTGGCCCGTTACCGCTGA
AAGGATATCCAACAGGTTGGAAACGCCCGCTTTGTTCTGCACATCGTAGCGAACTAC
CGGCGGCTCGTCGGAGTCAGTGACCGCACGTTTGATTTTCTTCACTACCGATTTCGG
ATCTTCCAGCAGGCCGATAACGTTATTGCGATTATCGTCAGACTTGGACATCTTCTTG
GTCGGCTCCAGCAGCGACATTACGCGCGCGCCAGATTTCGGAATAAACGGCTCCGG
CACCTTAAAGATCTCGCCATACAGCGCGTTGAAACGCTGGGCAATATCGCGGCTCA
GTTCGAGGTGCTGTTTCTGGTCTTCACCacaaggaccCAGATTAGTTTGATACAGCAGGA
TGTCCGCTGCCATCAGCACCGGATAGTCAAACAGACCAGCGTTGATGTTCTCGGCAT
AACGCGCAGATTTATCTTTAAACTGCGTCATGCGACTCAGTTCGCCGAAGTAGGTAT
AGCAGTTCAGTGCCCAGCCTAACTGTGCATGTTCCGGCACGTGGGACTGAACAAAA

ATGGTGCTTTTCTCAGGATCGATACCACAAGCCAGATACAAGGCCAGCGTATCCAGC
GTCGCTTTACGCAGCTTCTGTGCATCCTGGCGCACGGTGATCGCGTGTTGGTCAACG
ATACAGTAAATGCAATGGTAGTCATCCTGCATGTTTACCCACTGACGCAGCGCACCC
ATGTAGTTACCAATGGTCAATTCACCTGAGGGCTGTGCGCCagcAAAAACGATGGGCT
TAGTCATggtggcgctagccagcttgggtctccctatagtgagtcgtattaatttcgataagccagtaagcagtgggttctctagttagc
cagagagctctgcttatatagacctcccaccgtacacgcctaccgcccatttgcgtcaatggggcggagttgttacgacattttggaaagtcc
cgttgattttggtgccaaaacaaactcccattgacgtcaatggggtggagacttggaaatccccgtgagtcaaaccgctatccacgcccattg
atgtactgccaaaaccgcatcaccatggtaatagcgatgactaatacgtagatgtactgccaagtaggaaagtcccataaggtcatgtactgg
gcataatgccaggcgggccatttaccgtcattgacgtcaatagggggcgtacttggcatatgatacacttgatgtactgccaagtgggcagtt
taccgtaaatagtccacccattgacgtcaatggaaagtccctattggcgttactatgggaacatacgtcattattgacgtcaatgggcgggggt
cgttgggcggtcagccaggcgggccatttaccgtaagttatgtaacgcggaactccatatatgggctatgaactaatgaccccgtaattgatt
actattaataactagcaaaaaaTGGCAGGGGCGGAGAGACTCGAACTCCCAaCACCCGGTTTTgaA
GACCGGTGCTCTACCAATTGAACTACGCCCCTGGTGTTTCGTCCTTTCCACAAGATAT
ATAAAGCCAAGAAATCGAAATACTTTCAAGTTACGGTAAGCATATGATAGTCCATTT
TAAAACATAATTTTAAAACTGCAAACTACCCAAGAAATTATTACTTTCTACGTCACG
TATTTTGTACTAATATCTTTGTGTTTACAGTCAAATTAATTCTAATTATCTCTCTAACA
GCCTTGTATCGTATATGCAAATATGAAGGAATCATGGGAAATAGGCCCTCTTCCTGC
CCGACCTAGcaaaaaaTGGCAGGGGCGGAGAGACTCGAACTCCCAaCACCCGGTTTTgaA
GACCGGTGCTCTACCAATTGAACTACGCCCCTGGTGTTTCGTCCTTTCCACAAGATAT
ATAAAGCCAAGAAATCGAAATACTTTCAAGTTACGGTAAGCATATGATAGTCCATTT
TAAAACATAATTTTAAAACTGCAAACTACCCAAGAAATTATTACTTTCTACGTCACG
TATTTTGTACTAATATCTTTGTGTTTACAGTCAAATTAATTCTAATTATCTCTCTAACA
GCCTTGTATCGTATATGCAAATATGAAGGAATCATGGGAAATAGGCCCTCTTCCTGC
CCGACCTAGcaaaaaaTGGCAGGGGCGGAGAGACTCGAACTCCCAaCACCCGGTTTTgaA
GACCGGTGCTCTACCAATTGAACTACGCCCCTGGTGTTTCGTCCTTTCCACAAGATAT
ATAAAGCCAAGAAATCGAAATACTTTCAAGTTACGGTAAGCATATGATAGTCCATTT
TAAAACATAATTTTAAAACTGCAAACTACCCAAGAAATTATTACTTTCTACGTCACG
TATTTTGTACTAATATCTTTGTGTTTACAGTCAAATTAATTCTAATTATCTCTCTAACA
GCCTTGTATCGTATATGCAAATATGAAGGAATCATGGGAAATAGGCCCTCTTCCTGC
CCGACCTAGcaaaaaaTGGCAGGGGCGGAGAGACTCGAACTCCCAaCACCCGGTTTTgaA
GACCGGTGCTCTACCAATTGAACTACGCCCCTGGTGTTTCGTCCTTTCCACAAGATAT
ATAAAGCCAAGAAATCGAAATACTTTCAAGTTACGGTAAGCATATGATAGTCCATTT
TAAAACATAATTTTAAAACTGCAAACTACCCAAGAAATTATTACTTTCTACGTCACG
TATTTTGTACTAATATCTTTGTGTTTACAGTCAAATTAATTCTAATTATCTCTCTAACA
GCCTTGTATCGTATATGCAAATATGAAGGAATCATGGGAAATAGGCCCTCTTCCTGC
CCGACctagtcaataatcaatgtcaacgcgtatatctggcccgtacatcgcgaagcagcgcaaaacGGATCCtgcaggCTAG
TTATTAATAGTAATCAATTACGGGGTCATTAGTTCATAGCCCATATATGGAGTTCCG
CGTTACATAACTTACGGTAAATGGCCCGCCTGGCTGACCGCCCAACGACCCCCGCCC
ATTGACGTCAATAATGACGTATGTTCCCATAGTAACGCCAATAGGGACTTTCCATTG
ACGTCAATGGGTGGAgTATTTACGGTAAACTGCCCACTTGGCAGTACATCAAGTGTA
TCATATGCCAAGTACGCCCCCTATTGACGTCAATGACGGTAAATGGCCCGCCTGGCA
TTATGCCCAGTACATGACCTTATGGGACTTTCCTACTTGGCAGTACATCTACGTATTA
GTCATCGCTATTACCATGGTCGAGGTGAGCCCCACGTTCTGCTTCACTCTCCCCATCT
CCCCCCCCTCCCCACCCCCAATTTTGTATTTATTTATTTTTTAATTATTTTGTGCAGCG
ATGGGGGCGGGGGGGGGGGGGCGCGCGCCAGGCGGGGCGGGGCGGGGCGAGGG
GCGGGGCGGGGCGAGGCGGAGAGGTGCGGCGGCAGCCAATCAGAGCGGCGCGCTC

CGAAAGTTTCCTTTTATGGCGAGGCGGCGGCGGCGGCGGCCCTATAAAAAGCGAAG
CGCGCGGCGGGCGGGAGTCGCTGCGTTGCCTTCGCCCCGTGCCCCGCTCCGCGCCGC
CTCGCGCCGCCCGCCCCGGCTCTGACTGACCGCGTTACTCCCACAGGTGAGCGGGCG
GGACGGCCCTTCTCCTCCGGGCTGTAATTAGCGCTTGGTTTAATGACGGCTCGTTTCT
TTTCTGTGGCTGCGTGAAAGCCTTAAAGGGCTCCGGGAGGGCCCTTTGTGCGGGGGG
GAGCGGCTCGGGGGGTGCGTGCGTGTGTGTGTGCGTGGGGAGCGCCGCGTGCGGCC
CGCGCTGCCCGGCGGCTGTGAGCGCTGCGGGCGCGGCGCGGGGCTTTGTGCGCTCC
GCGTGTGCGCGAGGGGAGCGCGGCCGGGGGCGGTGCCCCGCGGTGCGGGGGGGCT
GCGAGGGGAACAAAGGCTGCGTGCGGGGTGTGTGCGTGGGGGGGTGAGCAGGGGG
TGTGGGCGCGGCGGTCGGGCTGTAACCCCCCCCTGCACCCCCCTCCCCGAGTTGCTG
AGCACGGCCCGGCTTCGGGTGCGGGGCTCCGTGCGGGGCGTGGCGCGGGGCTCGCC
GTGCCGGGCGGGGGGTGGCGGCAGGTGGGGGTGCCGGGCGGGGCGGGGCCGCCTC
GGGCCGGGGAGGGCTCGGGGGAGGGGCGCGGCGGCCCCGGAGCGCCGGCGGCTGT
CGAGGCGCGGCGAGCCGCAGCCATTGCCTTTTATGGTAATCGTGCGAGAGGGCGCA
GGGACTTCCTTTGTCCCAAATCTGGCGGAGCCGAAATCTGGGAGGCGCCGCCGCAC
CCCCTCTAGCGGGCGCGGGCGAAGCGGTGCGGCGCCGGCAGGAAGGAAATGGGCG
GGGAGGGCCTTCGTGCGTCGCCGCGCCGCCGTCCCCTTCTCCATCTCCAGCCTCGGG
GCTGCCGCAGGGGGACGGCTGCCTTCGGGGGGGACGGGGCAGGGCGGGGTTCGGCT
TCTGGCGTGTGACCGGCGGCTCTAGAGCCTCTGCTAACCATGTTCATGCCTTCTTCTT
TTTCCTACAGCTCCTGGGCAACGTGCTGGTTaTTGTGCTGTCTCATCATTTTGGCAAA
GAATTGGCCAAGGAGGCCACCATGGTGAGCAAGGGCGAGGAGCTGTTCACCGGGGT
GGTGCCCATCCTGGTCGAGCTGGACGGCGACGTAAACGGCCACAAGTTCAGCGTGT
CCGGCGAGGGCGAGGGCGATGCCACCTAgGGCAAGCTGACCCTGAAGTTCATCTGC
ACCACCGGCAAGCTGCCCGTGCCCTGGCCCACCCTCGTGACCACCCTGACCTACGGC
GTGCAGTGCTTCAGCCGCTACCCCGACCACATGAAGCAGCACGACTTCTTCAAGTCC
GCCATGCCCGAAGGCTACGTCCAGGAGCGCACCATCTTCTTCAAGGACGACGGCAA
CTACAAGACCCGCGCCGAGGTGAAGTTCGAGGGCGACACCCTGGTGAACCGCATCG
AGCTGAAGGGCATCGACTTCAAGGAGGACGGCAACATCCTGGGGCACAAGCTGGAG
TACAACTACAACAGCCACAACGTCTgaATCATGGCCGACAAGCAGAAGAACGGCATC
AAGGTGAACTTCAAGATCCGCCACAACATCGAGGACGGCAGCGTGCAGCTCGCCGA
CCACTACCAGCAGAACACCCCCATCGGCGACGGCCCCGTGCTGCTGCCCGACAACC
ACTACCTGAGCACCCAGTCCGCCCTGAGCAAAGACCCCAACGAGAAGCGCGATCAC
ATGGTCCTGCTGGAGTTCGTGACCGCCGCCGGGATCACTCTCGGCATGGACGAGCTG
TACAAGGGGCCCTTCGAACAAAAACTCATCTCAGAAGAGGATCTGAATATGCATAC
CGGTCATCATCACCATCACCATcatcaccatcaccatcatTaAggcctctaaggccgaattcaacgcgttaagtcgac
aatcaacctctggattacaaaatttgtgaaagattgactggtattcttaactatgttgctccttttacgctatgtggatacgctgctttaatgcctttgt
atcatgctattgcttcccgtatggctttcattttctcctccttgtataaatcctggttgctgtctctttatgaggagttgtggcccgttgtcaggcaac
gtggcgtggtgtgcactgtgtttgctgacgcaacccccactggttggggcattgccaccacctgtcagctcctttccgggactttcgctttccc
cctccctattgccacggcggaactcatcgccgcctgccttgcccgctgctggacaggggctcggctgttgggcactgacaattccgtggtgt
tgtcggggaaatcatcgtcctttccttggctgctcgcctgtgttgccacctggattctgcgcgggacgtccttctgctacgtcccttcggccctc
aatccagcggaccttccttcccgcggcctgctgccggctctgcggcctcttccgcgtcttcgccttcgccctcagacgagtcggatctcccttt
gggccgcctccccgcgtcgactttaactcggccagcacagtggtcgatcgaCCAATGCCCTGGCTCACAAATACC
ACTGAGATCTTTTTCCCTCTGCCAAAAATTATGGGGACATCATGAAGCCCCTTGAGC
ATCTGACTTCTGGCTAATAAAGGAAATTTATTTTCATTGCAATAGTGTGTTGGAATTT
TTTGTGTCTCTCACTCGGAAGGACATATGGGAGGGCAAATCATTTAAAACATCAGAA
TGAGTATTTGGTTTAGAGTTTGGCAACATATGCCcATATGCTGGCTGCCATGAACAA

AGGTtGGCTATAAAGAGGTCATCAGTATATGAAACAGCCCCCTGCTGTCCATTCCTTA
TTCCATAGAAAAGCCTTGACTTGAGGTTAGATTTTTTTTATATTTTGTTTTGTGTTATT
TTTTTCTTTAACATCCCTAAAATTTTCCTTACATGTTTTACTAGCCAGATTTTTCCTCC
TCTCCTGACTACTCCCAGTCATAGCTGTCCCTCTTCtttGCGGCCGCggtccgtatactccggaatat
taatagatcatggagataattaaaatgataaccatctcgcaaataaataagtattttactgttttcgtaacagttttgtaataaaaaaacctataaat
attccggattattcataccgtcccaccatcgggcgcgAACTCCTAAAAAACCGCCACCatgaagtgccttttgtacttagc
ctttttattcattggggtgaattgcaagttcaccatagtttttccacacaaccaaaaaggaaactggaaaaatgttccttctaattaccattattgcc
cgtcaagctcagatttaaattggcataatgacttaataggcacagccttacaagtcaaaatgcccaagagtcacaaggctattcaagcagacg
gttggatgtgtcatgcttccaaatgggtcactacttgtgatttccgctggtatggaccgaagtatataacacattccatccgatccttcactccatc
tgtagaacaatgcaaggaaagcattgaacaaacgaaacaaggaacttggctgaatccaggcttccctcctcaaagttgtggatatgcaactg
tgacggatgccgaagcagtgattgtccaggtgactcctcaccatgtgctggttgatgaatacacaggagaatgggttgattcacagttcatca
acggaaaatgcagcaattacatatgccccactgtccataactctacaacctggcattctgactataaggtcaaagggctatgtgattctaacct
catttccatggacatcaccttcttctcagaggacggagagctatcatccctgggaaaggagggcacagggttcagaagtaactactttgctta
tgaaactggaggcaaggcctgcaaaatgcaatactgcaagcattggggagtcagactcccatcaggtgtctggttcgagatggctgataag
gatctctttgctgcagccagattccctgaatgcccagaagggtcaagtatctctgctccatctcagacctcagtggatgtaagtctaattcagg
acgttgagaggatcttggattattccctctgccaagaaacctggagcaaaatcagagcgggtcttccaatctctccagtggatctcagctatct
tgctcctaaaaacccaggaaccggtcctgctttcaccataatcaatggtaccctaaaatactttgagaccagatacatcagagtcgatattgct
gctccaatcctctcaagaatggtcggaatgatcagtggaactaccacagaaagggaactgtgggatgactgggcaccatatgaagacgtg
gaaattggacccaatggagttctgaggaccagttcaggatataagtttcctttatacatgattggacatggtatgttggactccgatcttcatctta
gctcaaaggctcaggtgttcgaacatcctcacattcaagacgctgcttcgcaacttcctgatgatgagagtttattttttggtgatactgggctat
ccaaaaatccaatcgagcttgtagaaggttggttcagtagttggaaaagctctattgcctctttttctttatcatagggttaatcattggactattct
tggttctccgagttggtatccatctttgcattaaattaaagcacaccaagaaaagacagatttatacagacatagagatgaaccgacttggaaa
gtgataaAagcttgtcgagaagtactagaggatcataatcagccataccacatttgtagaggttttacttgctttaaaaaacctcccacacctc
cccctgaacctgaaacataaaatgaatgcaattgttgttgttaacttgtttattgcagcttataatggttacaaataaagcaatagcatcacaaattt
cacaaataaagcatttttttcactgcattctagttgtggtttgtccaaactcatcaatgtatcttatcatgtctggatctgatcactgcttgagcctag
gagatccgaaccagataagtgaaatctagttccaaactattttgtcatttttaattttcgtattagcttacgacgctacacccagttcccatctatttt
gtcactcttccctaaataatccttaaaaactccatttccacccctcccagttcccaactattttgtccgcccacagcggggcatttttcttcctgtta
tgtttttaatcaaacatcctgccaactccatgtgacaaaccgtcatcttcggctacttt

**pb3-OMeYRS-8xYtR-EGFP-10xHis-TEV-3xUAA:** EcYRS sequence is colored magenta, EGFP is colored green, His tag is colored purple, TEV cleavage site is colored orange, and TEV protease is colored blue.

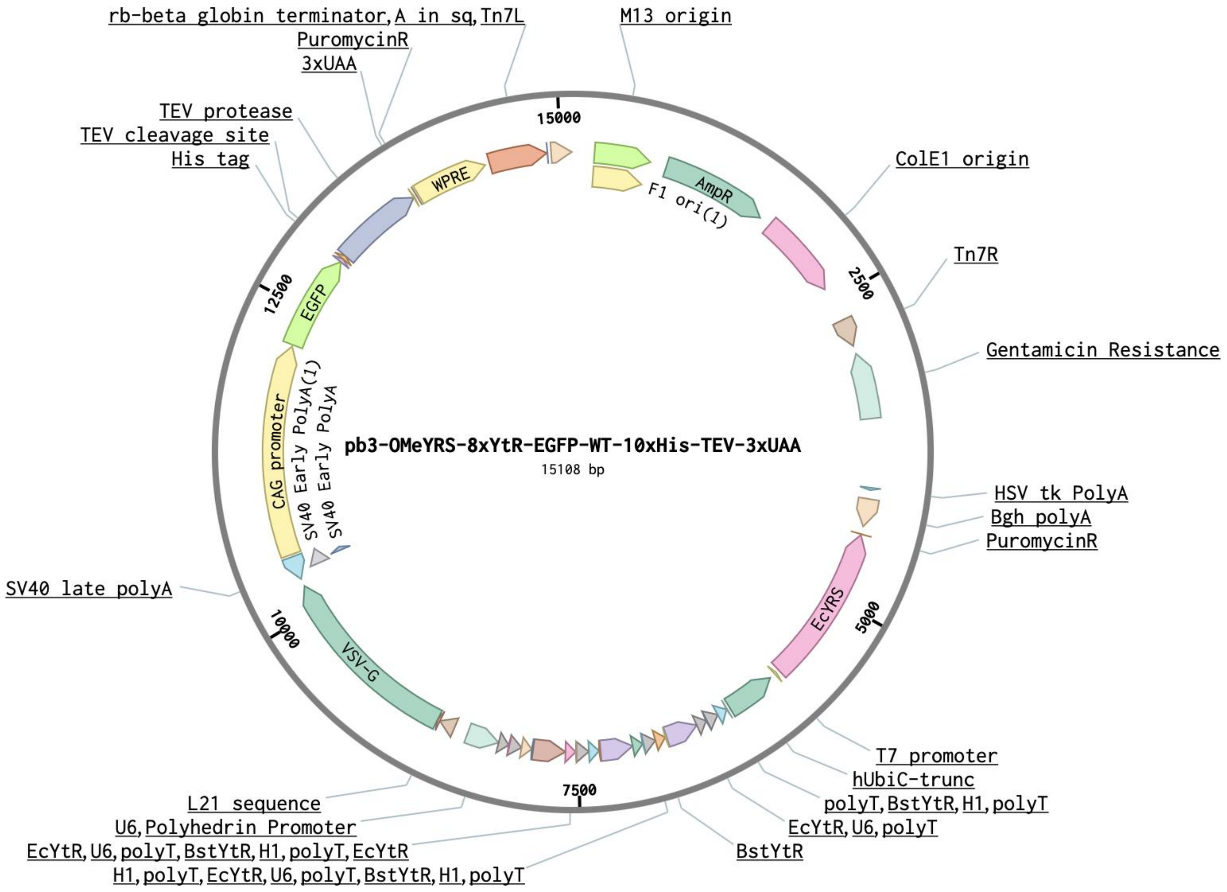


cctgttatgttttaatcaaacatcctgccaactccatgtgacaaaccgtcatcttcggctactttttctctgtcacagaatgaaatttttctgtcatc
tcttcgttattaatgtttgtaattgactgaatatcaacgcttatttgcagcctgaatggcgaatgggacgcgccctgtagcggcgcattaagcgc
ggcgggtgtggtggttacgcgcagcgtgaccgctacacttgccagcgccctagcgcccgctcctttcgctttcttcccttcctttctcgccacg
ttcgccggctttccccgtcaagctctaaatcgggggctccctttagggttccgatttagtgctttacggcacctcgaccccaaaaaacttgatta
gggtgatggttcacgtagtgggccatcgccctgatagacggtttttcgccctttgacgttggagtccacgttctttaatagtggactcttgttcca
aactggaacaacactcaaccctatctcggtctattcttttgatttataagggattttgccgatttcggcctattggttaaaaaatgagctgatttaac
aaaaatttaacgcgaattttaacaaaatattaacgtttacaatttcaggtggcacttttcggggaaatgtgcgcggaacccctatttgtttatttttct
aaatacattcaaatatgtatccgctcatgagacaataaccctgataaatgcttcaataatattgaaaaaggaagagtatgagtattcaacatttcc
gtgtcgcccttattcccttttttgcggcattttgccttcctgtttttgctcacccagaaacgctggtgaaagtaaaagatgctgaagatcagttggg
tgcacgagtgggttacatcgaactggatctcaacagcggtaagatccttgagagttttcgccccgaagaacgttttccaatgatgagcacttttt
aaagttctgctatgtggcgcggtattatcccgtattgacgccgggcaagagcaactcggtcgccgcatacactattctcagaatgacttggttg
agtactcaccagtcacagaaaagcatcttacggatggcatgacagtaagagaattatgcagtgctgccataaccatgagtgataacactgcg
gccaacttacttctgacaacgatcggaggaccgaaggagctaaccgcttttttgcacaacatgggggatcatgtaactcgccttgatcgttgg
gaaccggagctgaatgaagccataccaaacgacgagcgtgacaccacgatgcctgtagcaatggcaacaacgttgcgcaaactattaact
ggcgaactacttactctagcttcccggcaacaattaatagactggatggaggcggataaagttgcaggaccacttctgcgctcggcccttcc
ggctggctggtttattgctgataaatctggagccggtgagcgtgggtctcgcggtatcattgcagcactggggccagatggtaagccctccc

gtatcgtagttatctacacgacggggagtcaggcaactatggatgaacgaaatagacagatcgctgagataggtgcctcactgattaagcat
tggtaactgtcagaccaagtttactcatatatactttagattgatttaaaacttcatttttaatttaaaaggatctaggtgaagatcctttttgataatct
catgaccaaaatcccttaacgtgagttttcgttccactgagcgtcagaccccgtagaaaagatcaaaggatcttcttgagatcctttttttctgcg
cgtaatctgctgcttgcaaacaaaaaaaccaccgctaccagcggtggtttgtttgccggatcaagagctaccaactctttttccgaaggtaact
ggcttcagcagagcgcagataccaaatactgtccttctagtgtagccgtagttaggccaccacttcaagaactctgtagcaccgcctacatac
ctcgctctgctaatcctgttaccagtggctgctgccagtggcgataagtcgtgtcttaccgggttggactcaagacgatagttaccggataag
gcgcagcggtcgggctgaacggggggttcgtgcacacagcccagcttggagcgaacgacctacaccgaactgagatacctacagcgtg
agcattgagaaagcgccacgcttcccgaagggagaaaggcggacaggtatccggtaagcggcagggtcggaacaggagagcgcacg
agggagcttccagggggaaacgcctggtatctttatagtcctgtcgggtttcgccacctctgacttgagcgtcgatttttgtgatgctcgtcagg
ggggcggagcctatggaaaaacgccagcaacgcggcctttttacggttcctggccttttgctggccttttgctcacatgttctttcctgcgttat
cccctgattctgtggataaccgtattaccgcctttgagtgagctgataccgctcgccgcagccgaacgaccgagcgcagcgagtcagtgag
cgaggaagcggaagagcgcctgatgcggtattttctccttacgcatctgtgcggtatttcacaccgcagaccagccgcgtaacctggcaaa
atcggttacggttgagtaataaatggatgccctgcgtaagcgggtgtgggcggacaataaagtcttaaactgaacaaaatagatctaaactat
gacaataaagtcttaaactagacagaatagttgtaaactgaaatcagtccagttatgctgtgaaaaagcatactggacttttgttatggctaaag
caaactcttcattttctgaagtgcaaattgcccgtcgtattaaagaggggcgtggccaagggcatggtaaagactatattcgcggcgttgtga
caatttaccgaacaactccgcggccgggaagccgatctcggcttgaacgaattgttaggtggcggtacttgggtcgatatcaaagtgcatca
cttcttcccgtatgcccaactttgtatagagagccactgcgggatcgtcaccgtaatctgcttgcacgtagatcacataagcaccaagcgcgtt
ggcctcatgcttgaggagattgatgagcgcggtggcaatgccctgcctccggtgctcgccggagactgcgagatcatagatatagatctca
ctacgcggctgctcaaacctgggcagaacgtaagccgcgagagcgccaacaaccgcttcttggtcgaaggcagcaagcgcgatgaatgt
cttactacggagcaagttcccgaggtaatcggagtccggctgatgttgggagtaggtggctacgtctccgaactcacgaccgaaaagatca
agagcagcccgcatggatttgacttggtcagggccgagcctacatgtgcgaatgatgcccatacttgagccacctaactttgttttagggcga
ctgccctgctgcgtaacatcgttgctgctgcgtaacatcgttgctgctccataacatcaaacatcgacccacggcgtaacgcgcttgctgcttg
gatgcccgaggcatagactgtacaaaaaaacagtcataacaagccatgaaaaccgccactgcgccgttaccaccgctgcgttcggtcaag
gttctggaccagttgcgtgagcgcatacgctacttgcattacagtttacgaaccgaacaggcttatgtcaactgggttcgtgccttcatccgttt
ccacggtgtgcgtcacccggcaaccttgggcagcagcgaagtcgaggcatttctgtcctggctggcgaacgagcgcaaggtttcggtctc
cacgcatcgtcaggcattggcggccttgctgttcttctacggcaaggtgctgtgcacggatctgccctggcttcaggagatcggtagacctc
ggccgtcgcggcgcttgccggtggtgctgaccccggatgaagtggttcgcatcctcggttttctggaaggcgagcatcgtttgttcgcccag
gactctagctatagttctagtggttggctacgtacccgtagtggctatggcagggcttgcgcttaatgcgccgctacagggcgcgtggggat
accccctagagccccagctggttctttccgcctcagaagccatagagcccaccgcatccccagcatgcctgctattgtcttcccaatcctccc
ccttgctgtcctgccccaccccaccccccagaatagaatgacacctactcagacaatgcgatgcaatttcctcattttattaggaaaggacagt
gggagtggcaccttccagggtcaaggaaggcacgggggaggggcaaacaacagatggctggcaactagaaggcacagtcgaggctg
atcagcgggtttaaacgggccctctagactcgagttaaagtcgacttaacgcgttgaattcttaaacgggccctttccagcaaatcagacagt
aattctttttaccgcgacgcagtaaggtaaaacgaccaaacagacgatcttcttctttaaagaagtattcaggatcggactgtttttcaccgttaa
tggtgatggcattggaggcgatagttttacgtgcctgaccacgggaaggttgcagttcagaatcgaccagtgcctgcatcaggtctgcgccc
ttttccatctcaaccatcggtacgccgtcctgcgccagctgttcgaagtccgcttcactcagcgcactcaaagaaccgctgaacaggcattcg
gtaatacgttttgccgcctgtaaaccttcttcaccgtgaaccagacgagtcacctgctccgccagtacatactgggcgcgcggtgctttaccg
ctgtttttatcttcttcttccagggcgttgatctcttcaatgctcataaaggtgaagaacttcaggaagcggtaaacgtcggcatccgcagtgttg
atccagaactggtagaatttgtacgggctggttttcttcgggtccaaccagactgcgccgccttcagttttaccaaatttggtgccatctgcttta
gtgatcagcggaacggtcaggccaaacacctgattctgatgcagacgacgggtcaggtcgataccagaagtgatgttaccccattggtcag
aaccaccaatttgcagcaccacaccgtactgtttgttagcacaggccataccataaccctggagcaggttgtaggaaaactcagtgaacgaa
atcccctgaccttcacggttgagacgctgcttaaccgcttctttgttgatcatctggttaacggagaagtgtttgccaatatcgcgcaggaaggt
cagcacattcatattgccgaaccagtcataattattggccgcgatagcagagttttctccacagtcgaaatcgaggaacggggcaacctgctt
acggattttgtccacccactcctgaacagtttcttcggtgttcagcttacgctcggcagctttgaagctcgggtcgccaatcagacccgtcgcg
ccgcctaccagcgcaaccggcttgtggcccgcctgctggaagcgtttcaggcataacaatggaacaagatgccccaaatgcaagctgtca
gcggtaggatcgaagccacaaatgagtgcgatcgggccttgcgccagtcgctctgctaacgcttcctcgtccgtcacctgggctaccagcc
cccgctcttgcaattgtttaatcaagttactgcttgccatGGTGGCgctagccagcttgggtctccctatagtgagtcgtattaatttcgata

agccagtaagcagtgggttctctagttagccagagagctCtagaccaagtgacgatcacagcgatccacaaacaagaaccgcgacccaa
atcccggctgcgacggaactagctgtgccacacccggcgcgtccttatataatcatcggcgttcaccgccccacggagatccctccgcaga
atcgccgagaagggactacttttcctcgcctgttccgctctctggaaagaaaaccagtgccctagagtcacccaagtcccgtcctaaaatgtc
cttctgctgatactggggttctaaggccgagtcttatgagcagcgggccgctgtcctgagcgtccgggcggaaggatcaggacgctcgctg
cgcccttcgtctgacgtggcagcgctcgccgtgaggagggggcgcccgcgggaggcgccaaaacccggcgcggaggccttcgaac
ggCCACTAGCCCTAGCAAAAAATGGAGGGGGACGGATTCGAACCGCCGAACCCAAA
GGGAGCGGATTTAGAGTCCGCCGCGTTTAGCCACTTCGCTACCCCTCCGGTGTCTCT
ATCACTGATAGGGAACTTATAAGTCTCTATCACTGATAGGGATTTCACGTTTATGGT
GATTTCCCAGAACACATAGCGACATGCAAATATTAAAAAATGGTGGGGGAAGGATT
CGAACCTTCGAAGTCTGTGACGGCAGATTTAGAGTCTGCTCCCTTTGGCCGCTCGGG
AACCCCACCGGTGTTTCGTCCTTTCCACAAGATATATAAAGCCAAGAAATCGAAATA
CTTTCAAGTTACGGTAAGCATATGATAGTCCATTTTAAAACATAATTTTAAAACTGC
AAACTACCCAAGAAATTATTACTTTCTACGTCACGTATTTTGTACTAATATCTTTGTG
TTTACAGTCAAATTAATTCTAATTATCTCTCTAACAGCCTTGTATCGTATATGCAAAT
ATGAAGGAATCATGGGAAATAGGCCCTCTTCCTGCCCGAcCTAGCAAAAAATGGAG
GGGGACGGATTCGAACCGCCGAACCCAAAGGGAGCGGATTTAGAGTCCGCCGCGTT
TAGCCACTTCGCTACCCCTCCGGTGTCTCTATCACTGATAGGGAACTTATAAGTCTCT
ATCACTGATAGGGATTTCACGTTTATGGTGATTTCCCAGAACACATAGCGACATGCA
AATATTAAAAAATGGTGGGGGAAGGATTCGAACCTTCGAAGTCTGTGACGGCAGAT
TTAGAGTCTGCTCCCTTTGGCCGCTCGGGAACCCCACCGGTGTTTCGTCCTTTCCACA
AGATATATAAAGCCAAGAAATCGAAATACTTTCAAGTTACGGTAAGCATATGATAG
TCCATTTTAAAACATAATTTTAAAACTGCAAACTACCCAAGAAATTATTACTTTCTAC
GTCACGTATTTTGTACTAATATCTTTGTGTTTACAGTCAAATTAATTCTAATTATCTCT
CTAACAGCCTTGTATCGTATATGCAAATATGAAGGAATCATGGGAAATAGGCCCTCT
TCCTGCCCGACCTAGCAAAAAATGGAGGGGGACGGATTCGAACCGCCGAACCCAAA
GGGAGCGGATTTAGAGTCCGCCGCGTTTAGCCACTTCGCTACCCCTCCGGTGTCTCT
ATCACTGATAGGGAACTTATAAGTCTCTATCACTGATAGGGATTTCACGTTTATGGT
GATTTCCCAGAACACATAGCGACATGCAAATATTAAAAAATGGTGGGGGAAGGATT
CGAACCTTCGAAGTCTGTGACGGCAGATTTAGAGTCTGCTCCCTTTGGCCGCTCGGG
AACCCCACCGGTGTTTCGTCCTTTCCACAAGATATATAAAGCCAAGAAATCGAAATA
CTTTCAAGTTACGGTAAGCATATGATAGTCCATTTTAAAACATAATTTTAAAACTGC
AAACTACCCAAGAAATTATTACTTTCTACGTCACGTATTTTGTACTAATATCTTTGTG
TTTACAGTCAAATTAATTCTAATTATCTCTCTAACAGCCTTGTATCGTATATGCAAAT
ATGAAGGAATCATGGGAAATAGGCCCTCTTCCTGCCCGAcCTAGCAAAAAATGGAG
GGGGACGGATTCGAACCGCCGAACCCAAAGGGAGCGGATTTAGAGTCCGCCGCGTT
TAGCCACTTCGCTACCCCTCCGGTGTCTCTATCACTGATAGGGAACTTATAAGTCTCT
ATCACTGATAGGGATTTCACGTTTATGGTGATTTCCCAGAACACATAGCGACATGCA
AATATTAAAAAATGGTGGGGGAAGGATTCGAACCTTCGAAGTCTGTGACGGCAGAT
TTAGAGTCTGCTCCCTTTGGCCGCTCGGGAACCCCACCGGTGTTTCGTCCTTTCCACA
AGATATATAAAGCCAAGAAATCGAAATACTTTCAAGTTACGGTAAGCATATGATAG
TCCATTTTAAAACATAATTTTAAAACTGCAAACTACCCAAGAAATTATTACTTTCTAC
GTCACGTATTTTGTACTAATATCTTTGTGTTTACAGTCAAATTAATTCTAATTATCTCT
CTAACAGCCTTGTATCGTATATGCAAATATGAAGGAATCATGGGAAATAGGCCCTCT
TCCTGCCCGACctagtcaataatcaatgtcaacgcgtatatctggcccgtacatcgcgaagcagcgcaaaacGGATCCtgca
ggtatttGCGGCCGCggtccgtatactccggaatattaatagatcatggagataattaaaatgataaccatctcgcaaataaataagtatt
ttactgttttcgtaacagttttgtaataaaaaaacctataaatattccggattattcataccgtcccaccatcgggcgcgAACTCCTAAA

AAACCGCCACCatgaagtgccttttgtacttagcctttttattcattggggtgaattgcaagttcaccatagtttttccacacaaccaaa
aaggaaactggaaaaatgttccttctaattaccattattgcccgtcaagctcagatttaaattggcataatgacttaataggcacagccttacaa
gtcaaaatgcccaagagtcacaaggctattcaagcagacggttggatgtgtcatgcttccaaatgggtcactacttgtgatttccgctggtatg
gaccgaagtatataacacattccatccgatccttcactccatctgtagaacaatgcaaggaaagcattgaacaaacgaaacaaggaacttgg
ctgaatccaggcttccctcctcaaagttgtggatatgcaactgtgacggatgccgaagcagtgattgtccaggtgactcctcaccatgtgctg
gttgatgaatacacaggagaatgggttgattcacagttcatcaacggaaaatgcagcaattacatatgccccactgtccataactctacaacct
ggcattctgactataaggtcaaagggctatgtgattctaacctcatttccatggacatcaccttcttctcagaggacggagagctatcatccctg
ggaaaggagggcacagggttcagaagtaactactttgcttatgaaactggaggcaaggcctgcaaaatgcaatactgcaagcattgggga
gtcagactcccatcaggtgtctggttcgagatggctgataaggatctctttgctgcagccagattccctgaatgcccagaagggtcaagtatct
ctgctccatctcagacctcagtggatgtaagtctaattcaggacgttgagaggatcttggattattccctctgccaagaaacctggagcaaaat
cagagcgggtcttccaatctctccagtggatctcagctatcttgctcctaaaaacccaggaaccggtcctgctttcaccataatcaatggtacc
ctaaaatactttgagaccagatacatcagagtcgatattgctgctccaatcctctcaagaatggtcggaatgatcagtggaactaccacagaa
agggaactgtgggatgactgggcaccatatgaagacgtggaaattggacccaatggagttctgaggaccagttcaggatataagtttccttt
atacatgattggacatggtatgttggactccgatcttcatcttagctcaaaggctcaggtgttcgaacatcctcacattcaagacgctgcttcgc
aacttcctgatgatgagagtttatttttggtgatactgggctatccaaaaatccaatcgagcttgtagaaggttggttcagtagttggaaaagct
ctattgcctctttttctttatcatagggttaatcattggactattcttggttctccgagttggtatccatctttgcattaaattaaagcacaccaagaa
aagacagatttatacagacatagagatgaaccgacttggaaagtgataagtcgagaagtactagaggatcataatcagccataccacatttgt
agaggttttacttgctttaaaaaacctcccacacctccccctgaacctgaaacataaaatgaatgcaattgttgttgttaacttgtttattgcagctt
ataatggttacaaataaagcaatagcatcacaaatttcacaaataaagcatttttttcactgcattctagttgtggtttgtccaaactcatcaatgtat
cttatcatgtctggatctgatcactgcttgagcCTAGTTATTAATAGTAATCAATTACGGGGTCATTAGTT
CATAGCCCATATATGGAGTTCCGCGTTACATAACTTACGGTAAATGGCCCGCCTGGC
TGACCGCCCAACGACCCCCGCCCATTGACGTCAATAATGACGTATGTTCCCATAGTA
ACGCCAATAGGGACTTTCCATTGACGTCAATGGGTGGAgTATTTACGGTAAACTGCC
CACTTGGCAGTACATCAAGTGTATCATATGCCAAGTACGCCCCCTATTGACGTCAAT
GACGGTAAATGGCCCGCCTGGCATTATGCCCAGTACATGACCTTATGGGACTTTCCT
ACTTGGCAGTACATCTACGTATTAGTCATCGCTATTACCATGGTCGAGGTGAGCCCC
ACGTTCTGCTTCACTCTCCCCATCTCCCCCCCCTCCCCACCCCCAATTTTGTATTTATT
TATTTTTTAATTATTTTGTGCAGCGATGGGGGCGGGGGGGGGGGGGGCGCGCGCCA
GGCGGGGCGGGGCGGGGCGAGGGGCGGGGCGGGGCGAGGCGGAGAGGTGCGGCG
GCAGCCAATCAGAGCGGCGCGCTCCGAAAGTTTCCTTTTATGGCGAGGCGGCGGCG
GCGGCGGCCCTATAAAAAGCGAAGCGCGCGGCGGGCGGGAGTCGCTGCGTTGCCTT
CGCCCCGTGCCCCGCTCCGCGCCGCCTCGCGCCGCCCGCCCCGGCTCTGACTGACCG
CGTTACTCCCACAGGTGAGCGGGCGGGACGGCCCTTCTCCTCCGGGCTGTAATTAGC
GCTTGGTTTAATGACGGCTCGTTTCTTTTCTGTGGCTGCGTGAAAGCCTTAAAGGGCT
CCGGGAGGGCCCTTTGTGCGGGGGGGAGCGGCTCGGGGGGTGCGTGCGTGTGTGTG
TGCGTGGGGAGCGCCGCGTGCGGCCCGCGCTGCCCGGCGGCTGTGAGCGCTGCGGG
CGCGGCGCGGGGCTTTGTGCGCTCCGCGTGTGCGCGAGGGGAGCGCGGCCGGGGGC
GGTGCCCCGCGGTGCGGGGGGGCTGCGAGGGGAACAAAGGCTGCGTGCGGGGTGTG
TGCGTGGGGGGGTGAGCAGGGGGTGTGGGCGCGGCGGTCGGGCTGTAACCCCCCCC
TGCACCCCCCTCCCCGAGTTGCTGAGCACGGCCCGGCTTCGGGTGCGGGGCTCCGTG
CGGGGCGTGGCGCGGGGCTCGCCGTGCCGGGCGGGGGGTGGCGGCAGGTGGGGGT
GCCGGGCGGGGCGGGGCCGCCTCGGGCCGGGGAGGGCTCGGGGGAGGGGCGCGGC
GGCCCCGGAGCGCCGGCGGCTGTCGAGGCGCGGCGAGCCGCAGCCATTGCCTTTTA
TGGTAATCGTGCGAGAGGGCGCAGGGACTTCCTTTGTCCCAAATCTGGCGGAGCCG
AAATCTGGGAGGCGCCGCCGCACCCCCTCTAGCGGGCGCGGGCGAAGCGGTGCGGC
GCCGGCAGGAAGGAAATGGGCGGGGAGGGCCTTCGTGCGTCGCCGCGCCGCCGTCC

CCTTCTCCATCTCCAGCCTCGGGGCTGCCGCAGGGGGACGGCTGCCTTCGGGGGGA
CGGGGCAGGGCGGGGTTCGGCTTCTGGCGTGTGACCGGCGGCTCTAGAGCCTCTGCT
AACCATGTTCATGCCTTCTTCTTTTTCCTACAGCTCCTGGGCAACGTGCTGGTTaTTGT
GCTGTCTCATCATTTTGGCAAAGAATTGGCCAAGGAGGCCATGGTGAGCAAGGGCG
AGGAGCTGTTCACCGGGGTGGTGCCCATCCTGGTCGAGCTGGACGGCGACGTAAAC
GGCCACAAGTTCAGCGTGTCCGGCGAGGGCGAGGGCGATGCCACCTACGGCAAGCT
GACCCTGAAGTTCATCTGCACCACCGGCAAGCTGCCCGTGCCCTGGCCCACCCTCGT
GACCACCCTGACCTACGGCGTGCAGTGCTTCAGCCGCTACCCCGACCACATGAAGC
AGCACGACTTCTTCAAGTCCGCCATGCCCGAAGGCTACGTCCAGGAGCGCACCATCT
TCTTCAAGGACGACGGCAACTACAAGACCCGCGCCGAGGTGAAGTTCGAGGGCGAC
ACCCTGGTGAACCGCATCGAGCTGAAGGGCATCGACTTCAAGGAGGACGGCAACAT
CCTGGGGCACAAGCTGGAGTACAACTACAACAGCCACAACGTCTATATCATGGCCG
ACAAGCAGAAGAACGGCATCAAGGTGAACTTCAAGATCCGCCACAACATCGAGGAC
GGCAGCGTGCAGCTCGCCGACCACTACCAGCAGAACACCCCCATCGGCGACGGCCC
CGTGCTGCTGCCCGACAACCACTACCTGAGCACCCAGTCCGCCCTGAGCAAAGACC
CCAACGAGAAGCGCGATCACATGGTCCTGCTGGAGTTCGTGACCGCCGCCGGGATC
ACTCTCGGCATGGACGAGCTGTACAAGGGGCCCTTCGAACAAAAACTCATCTCAGA
AGAGGATCTGAATATGCATACCGGTcatcatcaccatcaccatcatcaccatcacgaaaatctttattttcaggtgga
ggaagtggagaaagTttgtttaaggggccgcgtgattacaacccgatatcgagcaccatttgtcatttgacgaatgaatctgatgggcacac
aacatcgttgtatggtattggatttggtcccttcatcattacaaacaagcacttgtttagaagaaataatggaacactgttggtccaatcactacat
ggtgtattcaaggtcaagaacaccacgactttgcaacaacacctcattgatgggagggacatgataattattcgcatgcctaaggatttccca
ccatttcctcaaaagctgaaatttagagagccacaaagggaagagcgcatatgtcttgtgacaaccaacttccaaactaagagcatgtctagc
atggtgtcagacactagttgcacattcccttcatctgatggcatattctggaagcattggattcaaaccaaggatgggcagtgtggcagtccat
tagtatcaactagagatgggttcattgttggtatacactcagcatcgaatttcaccaacacaaacaattatttcacaagcgtgccgaaaaacttc
atggaattgttgacaaatcaggaggcgcagcagtgggttagtggttggcgattaaatgctgactcagtattgtgggggggccataaagttttc
atggtgaaacctgaagagccttttcagccagttaaggaagcgactcaactcatgaatcgtcgtcgccgtcgcTAATAATAAggcct
ctaaggccgaattcaacgcgttaagtcgacaatcaacctctggattacaaaatttgtgaaagattgactggtattcttaactatgttgctcctttta
cgctatgtggatacgctgctttaatgcctttgtatcatgctattgcttcccgtatggctttcattttctcctccttgtataaatcctggttgctgtctcttt
atgaggagttgtggcccgttgtcaggcaacgtggcgtggtgtgcactgtgtttgctgacgcaacccccactggttggggcattgccaccacc
tgtcagctcctttccgggactttcgctttccccctccctattgccacggcggaactcatcgccgcctgccttgcccgctgctggacaggggct
cggctgttgggcactgacaattccgtggtgttgtcggggaaatcatcgtcctttccttggctgctcgcctgtgttgccacctggattctgcgcg
ggacgtccttctgctacgtcccttcggccctcaatccagcggaccttccttcccgcggcctgctgccggctctgcggcctcttccgcgtcttc
gccttcgccctcagacgagtcggatctccctttgggccgcctccccgcgtcgactttaactcggccagcacagtggtcgatcgaCCAAT
GCCCTGGCTCACAAATACCACTGAGATCTTTTTCCCTCTGCCAAAAATTATGGGGAC
ATCATGAAGCCCCTTGAGCATCTGACTTCTGGCTAATAAAGGAAATTTATTTTCATT
GCAATAGTGTGTTGGAATTTTTTGTGTCTCTCACTCGGAAGGACATATGGGAGGGCA
AATCATTTAAAACATCAGAATGAGTATTTGGTTTAGAGTTTGGCAACATATGCCcAT
ATGCTGGCTGCCATGAACAAAGGTtGGCTATAAAGAGGTCATCAGTATATGAAACAG
CCCCCTGCTGTCCATTCCTTATTCCATAGAAAAGCCTTGACTTGAGGTTAGATTTTTT
TTATATTTTGTTTTGTGTTATTTTTTTCTTTAACATCCCTAAAATTTTCCTTACATGTTT
TACTAGCCAGATTTTTCCTCCTCTCCTGACTACTCCCAGTCATAGCTGTCCCTCTTCT
CTTATGgAGATCCCTCGACCTGCcctaggagatccgaaccagataagtgaaatctagttccaaactattttgtcatttttaa
ttttcgtattagcttacgacgctacacccagttcccatctattttgtcactcttccctaaataatccttaaaaactccatttccacccctcccagttcc
caactattttgtccgcccacagcggggcatttttctt

pb3-pAAFRS-8xYtR-anti-Her2-HC121TAG-LC169TGA: pAAFRS sequence is colored blue, and the antibody expression cassette is colored in orange.

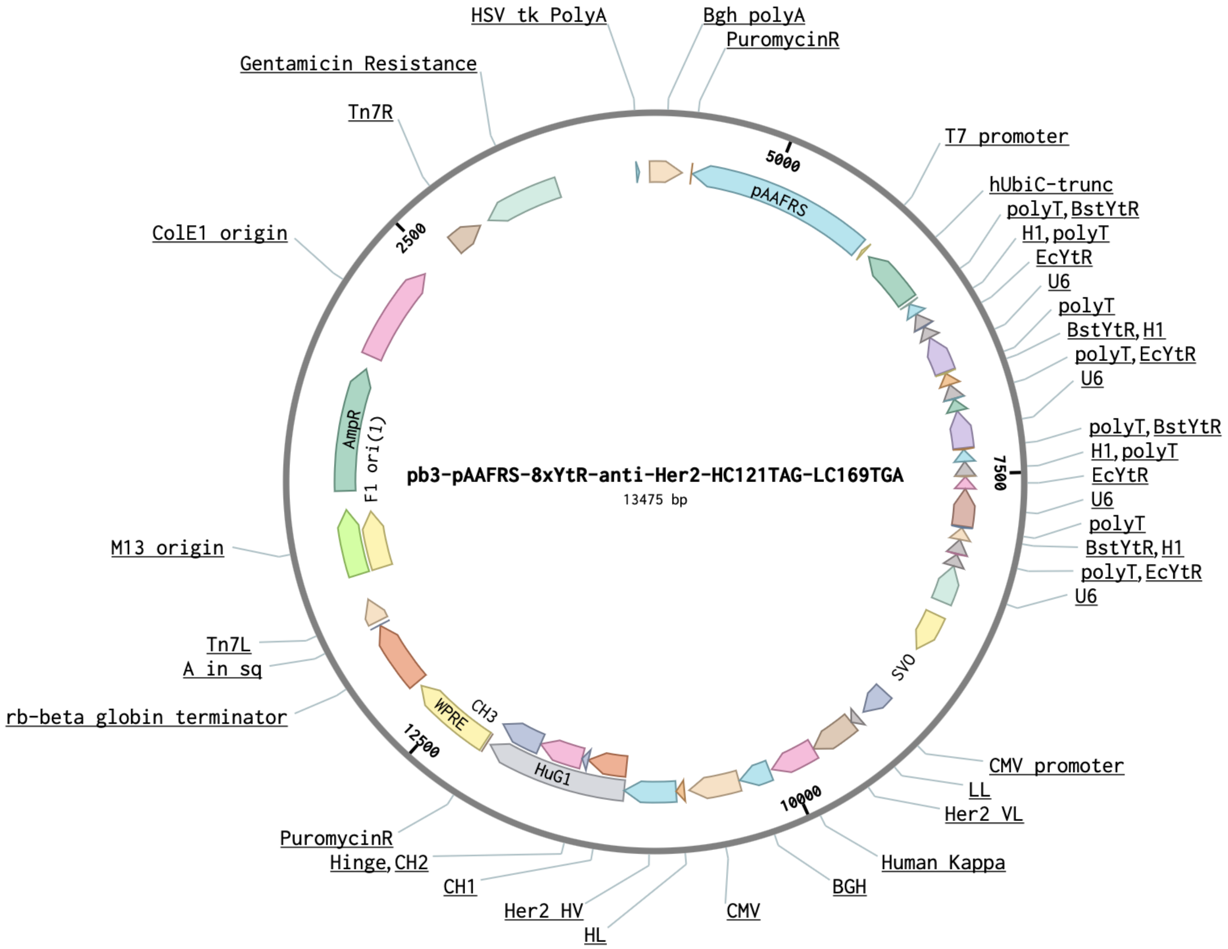


cctgttatgtttttaatcaaacatcctgccaactccatgtgacaaaccgtcatcttcggctactttttctctgtcacagaatgaaatttttctgtcatc
tcttcgttattaatgtttgtaattgactgaatatcaacgcttatttgcagcctgaatggcgaatgggacgcgccctgtagcggcgcattaagcgc
ggcgggtgtggtggttacgcgcagcgtgaccgctacacttgccagcgccctagcgcccgctcctttcgctttcttcccttcctttctcgccacg
ttcgccggctttccccgtcaagctctaaatcgggggctccctttagggttccgatttagtgctttacggcacctcgaccccaaaaaacttgatta
gggtgatggttcacgtagtgggccatcgccctgatagacggtttttcgccctttgacgttggagtccacgttctttaatagtggactcttgttcca
aactggaacaacactcaaccctatctcggtctattcttttgatttataagggattttgccgatttcggcctattggttaaaaaatgagctgatttaac
aaaaatttaacgcgaattttaacaaaatattaacgtttacaatttcaggtggcacttttcggggaaatgtgcgcggaacccctatttgtttatttttct
aaatacattcaaatatgtatccgctcatgagacaataaccctgataaatgcttcaataatattgaaaaaggaagagtatgagtattcaacatttcc
gtgtcgcccttattcccttttttgcggcattttgccttcctgtttttgctcacccagaaacgctggtgaaagtaaaagatgctgaagatcagttggg
tgcacgagtgggttacatcgaactggatctcaacagcggtaagatccttgagagttttcgccccgaagaacgttttccaatgatgagcactttt
aaagttctgctatgtggcgcggtattatcccgtattgacgccgggcaagagcaactcggtcgccgcatacactattctcagaatgacttggttg
agtactcaccagtcacagaaaagcatcttacggatggcatgacagtaagagaattatgcagtgctgccataaccatgagtgataacactgcg
gccaacttacttctgacaacgatcggaggaccgaaggagctaaccgcttttttgcacaacatgggggatcatgtaactcgccttgatcgttgg
gaaccggagctgaatgaagccataccaaacgacgagcgtgacaccacgatgcctgtagcaatggcaacaacgttgcgcaaactattaact

ggcgaactacttactctagcttcccggcaacaattaatagactggatggaggcggataaagttgcaggaccacttctgcgctcggcccttcc
ggctggctggtttattgctgataaatctggagccggtgagcgtgggtctcgcggtatcattgcagcactggggccagatggtaagccctccc
gtatcgtagttatctacacgacggggagtcaggcaactatggatgaacgaaatagacagatcgctgagataggtgcctcactgattaagcat
tggtaactgtcagaccaagtttactcatatatactttagattgatttaaaacttcatttttaatttaaaaggatctaggtgaagatcctttttgataatct
catgaccaaaatcccttaacgtgagttttcgttccactgagcgtcagaccccgtagaaaagatcaaaggatcttcttgagatcctttttttctgcg
cgtaatctgctgcttgcaaacaaaaaaaccaccgctaccagcggtggtttgtttgccggatcaagagctaccaactctttttccgaaggtaact
ggcttcagcagagcgcagataccaaatactgtccttctagtgtagccgtagttaggccaccacttcaagaactctgtagcaccgcctacatac
ctcgctctgctaatcctgttaccagtggctgctgccagtggcgataagtcgtgtcttaccgggttggactcaagacgatagttaccggataag
gcgcagcggtcgggctgaacggggggttcgtgcacacagcccagcttggagcgaacgacctacaccgaactgagatacctacagcgtg
agcattgagaaagcgccacgcttcccgaagggagaaaggcggacaggtatccggtaagcggcagggtcggaacaggagagcgcacg
agggagcttccagggggaaacgcctggtatctttatagtcctgtcgggtttcgccacctctgacttgagcgtcgatttttgtgatgctcgtcagg
ggggcggagcctatggaaaaacgccagcaacgcggcctttttacggttcctggccttttgctggccttttgctcacatgttctttcctgcgttat
cccctgattctgtggataaccgtattaccgcctttgagtgagctgataccgctcgccgcagccgaacgaccgagcgcagcgagtcagtgag
cgaggaagcggaagagcgcctgatgcggtattttctccttacgcatctgtgcggtatttcacaccgcagaccagccgcgtaacctggcaaa
atcggttacggttgagtaataaatggatgccctgcgtaagcgggtgtgggcggacaataaagtcttaaactgaacaaaatagatctaaactat
gacaataaagtcttaaactagacagaatagttgtaaactgaaatcagtccagttatgctgtgaaaaagcatactggacttttgttatggctaaag
caaactcttcattttctgaagtgcaaattgcccgtcgtattaaagaggggcgtggccaagggcatggtaaagactatattcgcggcgttgtga
caatttaccgaacaactccgcggccgggaagccgatctcggcttgaacgaattgttaggtggcggtacttgggtcgatatcaaagtgcatca
cttcttcccgtatgcccaactttgtatagagagccactgcgggatcgtcaccgtaatctgcttgcacgtagatcacataagcaccaagcgcgtt
ggcctcatgcttgaggagattgatgagcgcggtggcaatgccctgcctccggtgctcgccggagactgcgagatcatagatatagatctca
ctacgcggctgctcaaacctgggcagaacgtaagccgcgagagcgccaacaaccgcttcttggtcgaaggcagcaagcgcgatgaatgt
cttactacggagcaagttcccgaggtaatcggagtccggctgatgttgggagtaggtggctacgtctccgaactcacgaccgaaaagatca
agagcagcccgcatggatttgacttggtcagggccgagcctacatgtgcgaatgatgcccatacttgagccacctaactttgttttagggcga
ctgccctgctgcgtaacatcgttgctgctgcgtaacatcgttgctgctccataacatcaaacatcgacccacggcgtaacgcgcttgctgcttg
gatgcccgaggcatagactgtacaaaaaaacagtcataacaagccatgaaaaccgccactgcgccgttaccaccgctgcgttcggtcaag
gttctggaccagttgcgtgagcgcatacgctacttgcattacagtttacgaaccgaacaggcttatgtcaactgggttcgtgccttcatccgttt
ccacggtgtgcgtcacccggcaaccttgggcagcagcgaagtcgaggcatttctgtcctggctggcgaacgagcgcaaggtttcggtctc
cacgcatcgtcaggcattggcggccttgctgttcttctacggcaaggtgctgtgcacggatctgccctggcttcaggagatcggtagacctc
ggccgtcgcggcgcttgccggtggtgctgaccccggatgaagtggttcgcatcctcggttttctggaaggcgagcatcgtttgttcgcccag
gactctagctatagttctagtggttggctacgtacccgtagtggctatggcagggcttgcgcttaatgcgccgctacagggcgcgtggggat
acccccctagagccccagctggttctttccgcctcagaagccatagagcccaccgcatccccagcatgcctgctattgtcttcccaatcctccc
ccttgctgtcctgccccaccccaccccccagaatagaatgacacctactcagacaatgcgatgcaatttcctcattttattaggaaaggacagt
gggagtggcaccttccagggtcaaggaaggcacgggggaggggcaaacaacagatggctggcaactagaaggcacagtcgaggctg
atcagcgggtttaaacgggccctctagactcgagtttaaagtcgacttaacgcgttggaattcttatttccagcaaatcagacagtaattcttttta
ccgcgacgcagtaaggtaaaacgaccaaacagacgatcttcttctttaaagaagtattcaggatcggactgtttttcaccgttaatggtgatgg
cattggaggcgatagttttacgtgcctgaccacgggaaggttgcagttcagaatcgaccagtgcctgcatcaggtctgcgcccttttccatctc
aaccatcggtacgccgtcctgcgccagctgttcgaagtccgcttcactcagcgcactcaaagaaccgctgaacaggcattcggtaatacgtt
ttgccgcctgtaaaccttcttcaccgtgaaccagacgagtcacctgctccgccagtacatactgggcgcgcggtgctttaccgctgtttttatct
tcttcttccagggcgttgatctcttcaatgctcataaaggtgaagaacttcaggaagcggtaaacgtcggcACGcgcagtgttgatccaga
actggtagaatttgtacgggctggttttcttcggatcTaaccagactgcgccgccttcagttttaccaaatttggtgccatctgctttagtgatca
gcggaacggtcaggccaaacacctgattctgatgcagacgacgggtcaggtcgataccagaagtgatgttaccccactggtcagaaccac
caatttgcagcaccacaccgtactgtttgttACAacaggcATAACAataaccctgCaAcaggttgtaggaaaactcagtgaacga
aatcccctgaTcttcacggttgagacgctgcttaaccgcttctttgttgatcatctggttaacggagaagtgtttgccaatatcgcgcaggaag
gtcagcacattcatattgccgaaccagtcataGttGttCgccgcgatagcagagttttctccacagtcgaaatcgaggaacggggcaacct
gcttacggattttgtccacccactcctgaacagtttcttcggtgttcagcttacgctcggcagctttgaagctcgggtcgccaatcagacccgtc
gcgccgcctacAACcgcaaccggcttgtggcccgcctgctggaagcgtttcaggcataacaatggaacaagatgccccaatgcaagc

tgtcagcggtaggatcgaagccGcaCCCgagcgcgatcgggccttgcgccagtcgctctgctaacgcttcctcgtccgtcacctgggct
accagcccccgctcttgcaattgtttaatcaagttactgcttgccatGGTGGCgctagctagccagcttgggtctccctatagtgagtcgt
attaatttcgataagccagtaagcagtgggttctctagttagccagagagctCtagaccaagtgacgatcacagcgatccacaaacaagaa
ccgcgacccaaatcccggctgcgacggaactagctgtgccacacccggcgcgtccttatataatcatcggcgttcaccgccccacggaga
tccctccgcagaatcgccgagaagggactacttttcctcgcctgttccgctctctggaaagaaaaccagtgccctagagtcacccaagtccc
gtcctaaaatgtccttctgctgatactggggttctaaggccgagtcttatgagcagcgggccgctgtcctgagcgtccgggcggaaggatca
ggacgctcgctgcgcccttcgtctgacgtggcagcgctcgccgtgaggagggggcgcccgcgggaggcgccaaacccggcgcgg
aggccttcgaacggCCACTAGCCCTAGCAAAAAATGGAGGGGGACGGATTCGAACCGCCGA
ACCCAAAGGGAGCGGATTTAGAGTCCGCCGCGTTTAGCCACTTCGCTACCCCTCCGG
TGTCTCTATCACTGATAGGGAACTTATAAGTCTCTATCACTGATAGGGATTTCACGTT
TATGGTGATTTCCCAGAACACATAGCGACATGCAAATATTAAAAAATGGTGGGGGA
AGGATTCGAACCTTCGAAGTCTGTGACGGCAGATTTAGAGTCTGCTCCCTTTGGCCG
CTCGGGAACCCCACCGGTGTTTCGTCCTTTCCACAAGATATATAAAGCCAAGAAATC
GAAATACTTTCAAGTTACGGTAAGCATATGATAGTCCATTTTAAAACATAATTTTAA
AACTGCAAACTACCCAAGAAATTATTACTTTCTACGTCACGTATTTTGTACTAATATC
TTTGTGTTTACAGTCAAATTAATTCTAATTATCTCTCTAACAGCCTTGTATCGTATAT
GCAAATATGAAGGAATCATGGGAAATAGGCCCTCTTCCTGCCCGAcCTAGCAAAAA
ATGGAGGGGGACGGATTCGAACCGCCGAACCCAAAGGGAGCGGATTTAGAGTCCGC
CGCGTTTAGCCACTTCGCTACCCCTCCGGTGTCTCTATCACTGATAGGGAACTTATAA
GTCTCTATCACTGATAGGGATTTCACGTTTATGGTGATTTCCCAGAACACATAGCGA
CATGCAAATATTAAAAAATGGTGGGGGAAGGATTCGAACCTTCGAAGTCTGTGACG
GCAGATTTAGAGTCTGCTCCCTTTGGCCGCTCGGGAACCCCACCGGTGTTTCGTCCTT
TCCACAAGATATATAAAGCCAAGAAATCGAAATACTTTCAAGTTACGGTAAGCATA
TGATAGTCCATTTTAAAACATAATTTTAAAACTGCAAACTACCCAAGAAATTATTAC
TTTCTACGTCACGTATTTTGTACTAATATCTTTGTGTTTACAGTCAAATTAATTCTAAT
TATCTCTCTAACAGCCTTGTATCGTATATGCAAATATGAAGGAATCATGGGAAATAG
GCCCTCTTCCTGCCCGACCTAGCAAAAAATGGAGGGGGACGGATTCGAACCGCCGA
ACCCAAAGGGAGCGGATTTAGAGTCCGCCGCGTTTAGCCACTTCGCTACCCCTCCGG
TGTCTCTATCACTGATAGGGAACTTATAAGTCTCTATCACTGATAGGGATTTCACGTT
TATGGTGATTTCCCAGAACACATAGCGACATGCAAATATTAAAAAATGGTGGGGGA
AGGATTCGAACCTTCGAAGTCTGTGACGGCAGATTTAGAGTCTGCTCCCTTTGGCCG
CTCGGGAACCCCACCGGTGTTTCGTCCTTTCCACAAGATATATAAAGCCAAGAAATC
GAAATACTTTCAAGTTACGGTAAGCATATGATAGTCCATTTTAAAACATAATTTTAA
AACTGCAAACTACCCAAGAAATTATTACTTTCTACGTCACGTATTTTGTACTAATATC
TTTGTGTTTACAGTCAAATTAATTCTAATTATCTCTCTAACAGCCTTGTATCGTATAT
GCAAATATGAAGGAATCATGGGAAATAGGCCCTCTTCCTGCCCGAcCTAGCAAAAA
ATGGAGGGGGACGGATTCGAACCGCCGAACCCAAAGGGAGCGGATTTAGAGTCCGC
CGCGTTTAGCCACTTCGCTACCCCTCCGGTGTCTCTATCACTGATAGGGAACTTATAA
GTCTCTATCACTGATAGGGATTTCACGTTTATGGTGATTTCCCAGAACACATAGCGA
CATGCAAATATTAAAAAATGGTGGGGGAAGGATTCGAACCTTCGAAGTCTGTGACG
GCAGATTTAGAGTCTGCTCCCTTTGGCCGCTCGGGAACCCCACCGGTGTTTCGTCCTT
TCCACAAGATATATAAAGCCAAGAAATCGAAATACTTTCAAGTTACGGTAAGCATA
TGATAGTCCATTTTAAAACATAATTTTAAAACTGCAAACTACCCAAGAAATTATTAC
TTTCTACGTCACGTATTTTGTACTAATATCTTTGTGTTTACAGTCAAATTAATTCTAAT
TATCTCTCTAACAGCCTTGTATCGTATATGCAAATATGAAGGAATCATGGGAAATAG
GCCCTCTTCCTGCCCGACctagtcaataatcaatgtcaacgcgtatatctggcccgtacatcgcgaagcagcgcaaaacG

GATCCcgcggtgagttcaggcttaattaagtacgggcctccaaaaaagcctcctcactacttctggaatagctcagaggcagaggcgg
cctcggcctctgcataaataaaaaaaattagtcagccatggggcggagaatgggcggaactgggcggagttaggggcgggatgggcgg
agttaggggcgggactatggttgctgactaattgagatgcatgctttgcatacttctgcctgctggggagcctggggactttccacacctggtt
gctgactaattgagatgcatgctttgcatacttctgcccgcggagttattaatagtaatcaattacggggtcattagttcatagcccatatatgga
gttccgcgttacataacttacggtaaatggcccgcctggctgaccgcccaacgacccccgcccattgacgtcaataatgacgtatgttcccat
agtaacgccaatagggactttccattgacgtcaatgggtggagtatttacggtaaactgcccacttggcagtacatcaagtgtatcatatgcca
agtacgccccctattgacgtcaatgacggtaaatggcccgcctggcattatgcccagtacatgaccttatgggactttcctacttggcagtaca
tctacgtattagtcatcgctattaccatggtgatgcggttttggcagtacatcaatgggcgtggatagcggtttgactcacggggatttccaagt
ctccaccccattgacgtcaatgggagtttgttttggcaccaaaatcaacgggactttccaaaatgtcgtaacaactccgccccattgacgcaa
atgggcggtaggcgtgtacggtgggaggtctatataagcagagctgggtacgtgaaccgtcagatcgcctggagacgccatcacactagt
caccatgagggtccccgctcagctcctggggctcctgctgctctggctcccaggtgcacgatgtgacatccagatgacccagtccccctcct
ccctgtctgcctccgtgggcgacagagtgaccatcacctgtcgggcctcccaggatgtgaacaccgccgtggcctggtatcagcagaagc
ctggcaaggcccctaagctgctgatctactccgcctccttcctgtactccggcgtgccctcccggttctccggctccagatccggcaccgact
tcaccctgaccatctccagcctgcagcctgaggacttcgccacctactactgccagcagcactacaccacccctccaaccttcggccaggg
caccaaggtggagatcaagcgtacggtggctgcaccatctgtcttcatcttcccgccatctgatgagcagttgaaatctggaactgcctctgtt
gtgtgcctgctgaataacttctatcccagagaggccaaagtacagtggaaggtggataacgccctccaatcgggtaactcccaggagagtg
tcacagagcaggacagctgagacagcacctacagcctcagcagcaccctgacgctgagcaaagcagactacgagaaacacaaagtcta
cgcctgcgaagtcacccatcagggcctgagctcgcccgtcacaaagagcttcaacaggggagagtgttaataacctgcaggtcactgtgc
cttctagttgccagccatctgttgtttgcccctcccccgtgccttccttgaccctggaaggtgccactcccactgtcctttcctaataaaatgagg
aaattgcatcgcattgtctgagtaggtgtcattctattctggggggtggggtggggcaggacagcaagggggaggattgggaagacaatag
caggcatgctggggatgcggtggggtcgacatatgccaagtacgccccctattgacgtcaatgacggtaaatggcccgcctggcattatgc
ccagtacatgaccttatgggactttcctacttggcagtacatctacgtattagtcatcgctattaccatggtgatgcggttttggcagtacatcaat
gggcgtggatagcggtttgactcacggggatttccaagcctccaccccattgacgtcaatgggagtttgttttggcaccaaaatcaacggga
ctttccaaaatgtcgtaacaactccgccccattgacgcaaatgggcggtaggcgtgtacggtgggaggtctatataagcagagctgggtac
gtcctcacattcagtgatcagcactgaacacaggatatccaccatgggttggagcctcatcttgctcttccttgtcgctgttgctacgcgtgtcc
actccgaagtgcagctggtggagtctggcggaggactggtgcagccagggggcagcctgagactgtcttgcgccgcctccggcttcaac
atcaaggacacctacatccactgggtccgccaggcaccaggcaagggactggaatgggtggcccggatctaccctaccaacggctacac
cagatacgccgactccgtgaagggccggttcaccatctccgccgacacctccaagaacaccgcctacctgcagatgaattccctgagggc
cgaggacaccgccgtgtactactgctccagatggggaggcgacggcttctacgccatggactactggggccagggcaccctggtcacag
tgtcctcttagagcaccaagggcccatcggtcttccccctggcaccctcctccaagagcacctctgggggcacagcggccctgggctgcct
ggtcaaggactacttccccgaaccggtgacggtgtcgtggaactcaggcgccctgaccagcggcgtgcacaccttcccggctgtcctaca
gtcctcaggactctactccctcagcagcgtggtgaccgtgccctccagcagcttgggcacccagacctacatctgcaacgtgaatcacaag
cccagcaacaccaaggtggacaagaaagttgagcccaaatcttgtgacaaaactcacacatgcccaccgtgcccagcacctgaactcctg
gggggaccgtcagtcttcctcttccccccaaaacccaaggacaccctcatgatctcccggacccctgaggtcacatgcgtggtggtggacg
tgagccacgaagaccctgaggtcaagttcaactggtacgtggacggcgtggaggtgcataatgccaagacaaagccgcgggaggagca
gtacaacagcacgtaccgtgtggtcagcgtcctcaccgtcctgcaccaggactggctgaatggcaaggagtacaagtgcaaggtctccaa
caaagccctcccagcccccatcgagaaaaccatctccaaagccaaagggcagccccgagaaccacaggtgtacaccctgcccccatcc
cgggatgagctgaccaagaaccaggtcagcctgacctgcctggtcaaaggcttctatcccagcgacatcgccgtggagtgggagagcaa
tgggcagccggagaacaactacaagaccacgcctcccgtgctggactccgacggctccttcttcctctacagcaagctcaccgtggacaa
gagcaggtggcagcaggggaacgtcttctcatgctccgtgatgcatgaggctctgcacaaccactacacgcagaagagcctctccctgtct
ccgggtTCTtaaggcctctaaggccgaattcaacgcgttaagtcgacaatcaacctctggattacaaaatttgtgaaagattgactggtattc
ttaactatgttgctccttttacgctatgtggatacgctgctttaatgcctttgtatcatgctattgcttcccgtatggctttcattttctcctccttgtataa
atcctggttgctgtctctttatgaggagttgtggcccgttgtcaggcaacgtggcgtggtgtgcactgtgtttgctgacgcaacccccactggtt
ggggcattgccaccacctgtcagctcctttccgggactttcgctttccccctccctattgccacggcggaactcatcgccgcctgccttgccc
gctgctggacaggggctcggctgttgggcactgacaattccgtggtgttgtcggggaaatcatcgtcctttccttggctgctcgcctgtgttgc
cacctggattctgcgcgggacgtccttctgctacgtcccttcggccctcaatccagcggaccttccttcccgcggcctgctgccggctctgcg

gcctcttccgcgtcttcgccttcgccctcagacgagtcggatctccctttgggccgcctccccgcgtcgactttaactcggccagcacagtgg
tcgatcgaCCAATGCCCTGGCTCACAAATACCACTGAGATCTTTTTCCCTCTGCCAAAAA
TTATGGGGACATCATGAAGCCCCTTGAGCATCTGACTTCTGGCTAATAAAGGAAATT
TATTTTCATTGCAATAGTGTGTTGGAATTTTTTGTGTCTCTCACTCGGAAGGACATAT
GGGAGGGCAAATCATTTAAAACATCAGAATGAGTATTTGGTTTAGAGTTTGGCAAC
ATATGCCcATATGCTGGCTGCCATGAACAAAGGTtGGCTATAAAGAGGTCATCAGTA
TATGAAACAGCCCCCTGCTGTCCATTCCTTATTCCATAGAAAAGCCTTGACTTGAGG
TTAGATTTTTTTTATATTTTGTTTTGTGTTATTTTTTTCTTTAACATCCCTAAAATTTTC
CTTACATGTTTTACTAGCCAGATTTTTCCTCCTCTCCTGACTACTCCCAGTCATAGCT
GTCCCTCTTCTCTTATGgAGATCCCTCGACCTGCcctaggagatccgaaccagataagtgaaatctagttcca
aactattttgtcatttttaattttcgtattagcttacgacgctacacccagttcccatctattttgtcactcttccctaaataatccttaaaaactccattt
ccacccctcccagttcccaactattttgtccgcccacagcggggcatttttctt
